# A stepwise, modular design of building uniform brain assembloids representing the dynamic cellular interplay between neurons and glial cells

**DOI:** 10.1101/2024.10.01.615952

**Authors:** Eunjee Kim, Yunhee Kim, Soojung Hong, Inha Kim, Juhee Lee, Jong-Yeon Yoo, Jihyun Kim, Kwangmin Yoo, Joung-Hun Kim, Jungmin Choi, Kunyoo Shin

**Affiliations:** Institute of Molecular Biology and Genetics, Seoul National University, Seoul 08826, Republic of Korea; School of Biological Sciences, College of Natural Sciences, Seoul National University, Seoul 08826, Republic of Korea; Department of Life Sciences, Pohang University of Science and Technology, Pohang, Gyeongbuk 37673, Republic of Korea; Department of Biomedical Sciences, Korea University College of Medicine, Seoul 02841, Republic of Korea

**Keywords:** Brain assembloid, brain organoid, human brain development

## Abstract

Current brain organoid technology fails to provide adequate patterning cues to induce a mature structure that represent the complexity of the human brain. Here, we developed a module-based cellular reconstitution technology to sequentially build uniform forebrain assembloids with mature cortical structures and functional connectivity. The uniformity and maturity of the newly-conceived forebrain assembloids were achieved by creating single-rosette-based organoids at the early stage, whose sizes were big and consistent with the treatment of Wnt and Hedgehog agonists, followed by spatial reconstitution with the Reelin-expressing neuronal layer and non-neuronal glial cells. The resulting single-rosette-based forebrain assembloids were highly uniform and reproducible without significant batch effects, solving major heterogeneity issues caused by difficulties in controlling the number and size of rosettes in conventional multi-rosette organoids. Furthermore, these forebrain assembloids structurally and functionally recapitulated the physiology of the human brain, including the six-layered cortical structure, functional connectivity, and dynamic cellular interplay between neurons and glial cells. Our study thus provided an innovative preclinical model to study a range of neurological disorders, understanding the pathogenesis of which requires an organoid system capable of representing the dynamic cellular interactions and the maturity of the human brain.

## Introduction

Brain organoids are three-dimensional (3D), self-organizing tissues derived from pluripotent stem cells that have been developed for several years, starting from whole-brain organoids ^1,2^ to region-specific organoids—such as cortical organoids ^3–5^, midbrain organoids ^3,6,7^, and subpallial organoids ^4,8,9^—to connected hybrid organoids ^4,10,11^. Unlike conventional neural cell cultures and genetically-modified mouse models, which are not likely to represent the complex dynamics of the cellular interactions present in the human brain, brain organoids mimic many aspects of the tissue architecture and cellular composition of the human brain *in vivo*, thus serving as an innovative platform for studying the pathophysiology of the human brain ^12–14^.

Although brain organoids provide a valuable system for studying the human brain, current brain organoids have several critical limitations. First, the heterogeneity in structural and functional maturity between individual organoids and batches are problematic. The reproducibility issue in current forebrain organoid technologies hinders the rapid advance of this field, mainly due to the challenge of precisely controlling the quality of organoid growth during extended culture periods, which can last up to one year. Particularly, there is significant difficulty in the early control of the number and the size of vesicle-like, neuroepithelial structures called rosettes in conventional multi-rosette organoids. These structures are different from the secondary vesicle of telencephalon^15^ in *in vivo* developing human brain, which contains a single ventricular space and becomes forebrain^16^. Additionally, there is a lack of spatial and temporal control of reelin (RELN), which is expressed in the first layer of neurons, migrating into the marginal zone at the early stage of human corticogenesis, and is required for proper neural guidance and laminar organization^17,18^. Although several approaches have been used to increase the maturity of brain organoids, such as employing long-term culture methods^19,20^, these organoids still exhibit inconsistency and heterogeneity in their structural and functional maturity due to the lack of quality controls on organoid growth during extended culture periods. Second, currently available brain organoids show a low degree of cellular diversity. Despite several methodologies of co-culturing with glial cells, these models still exhibit immature cellular interactions between neurons and glial cells with a high level of inconsistency^21,22^. These limitations further hinder the study of dynamic cellular interactions in the human brain, rendering them incapable of examining cell-type-specific phenotypes and accurately modeling various disease-specific human brains. Therefore, combined with inconsistent development of structural and functional maturity and a lack of cellular diversity, current brain organoid technologies face substantial challenges that impede their ability to practically and reliably model the complexity of the human brain.

In this study, we developed a step-wise, module-based cellular reconstitution technology to sequentially build human forebrain assembloids ^23^. These single-rosette-based forebrain assembloids, developed by sequential reconstitution with non-neuronal glial cells, showed a great degree of consistency and uniformity with minimal heterogeneity and batch effects. Also, these forebrain assembloids represented the six-layered cortical structure with mature laminar organization and functional connectivity between neurons and glial cells.

## Results

### Creation of forebrain assembloids by a module-based, cellular reconstitution of multiple cell types in human brains

Current brain organoids exhibit several immature characteristics, including the lack of sustained proliferation of neural progenitor cells (NPCs), disorganized or absent populations of late-born neurons (layers 2–4), spatial and temporal inconsistency of the RELN-expressing first layer of neurons (Cajal–Retzius cells) for proper neural guidance and laminar organization, as well as a low degree of cellular diversity and a high level of heterogeneity and inconsistency between organoids ^3–5^. To overcome these limitations, we developed a stepwise strategy to create mature forebrain assembloids ^23^, aimed at providing essential developmental cues and reconstituting diverse different cell types within the system (Fig. 1a).

**Fig. 1.**
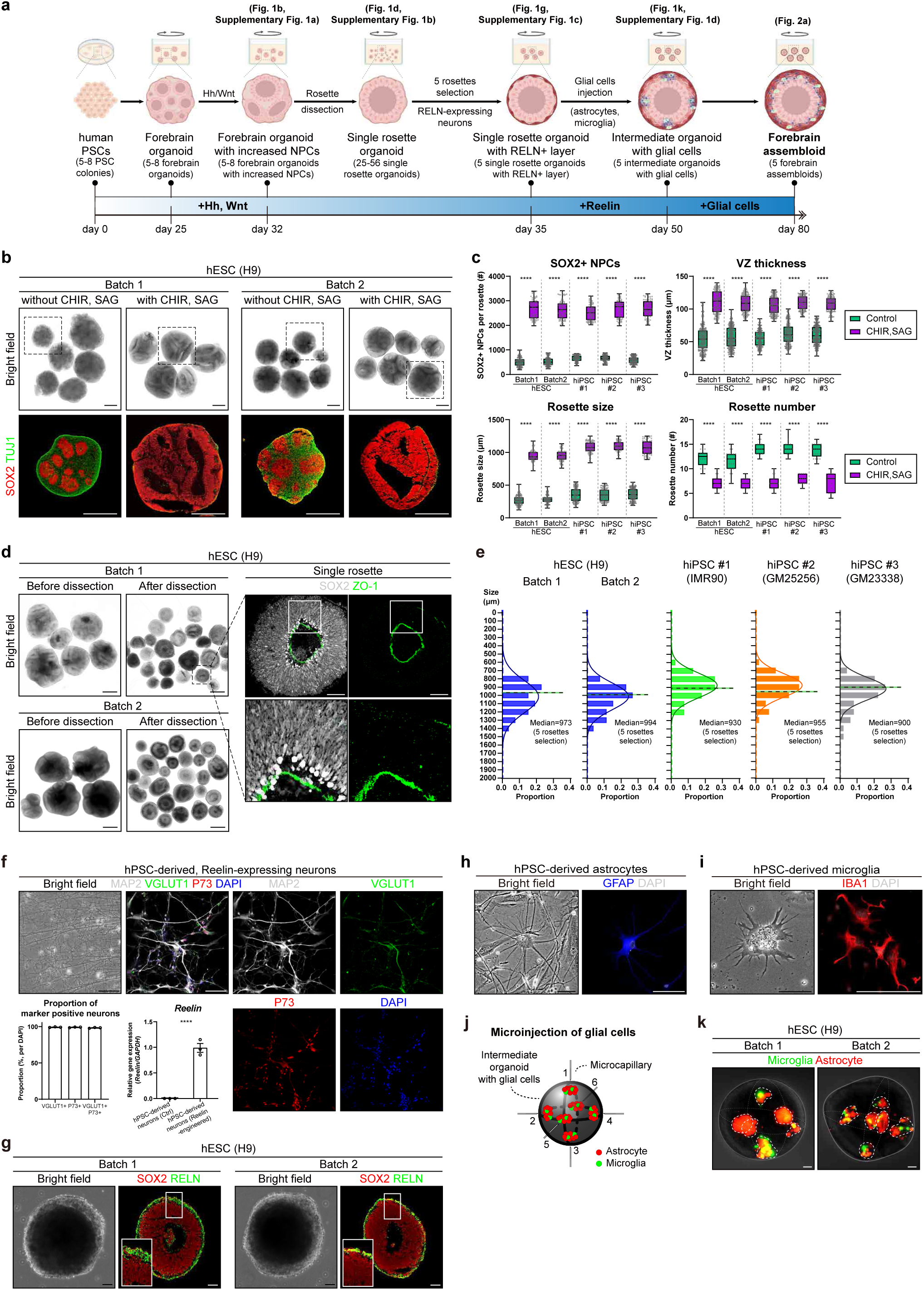
A stepwise, modular design of building uniform human forebrain assembloids. (a) Experimental scheme of creating forebrain assembloids. (b) (top) Representative bright field images of early forebrain organoids (day 32), derived from hESCs, treated with CHIR99021 and SAG. (bottom) Magnified images of forebrain organoids, demarcated by dotted boxes on top panels, immunostained for NPCs (SOX2, red) and neurons (TUJ1, green). Scale bars, 1 mm. (c) (top left) Quantification of the NPC population by counting the number of SOX2-positive cells in each rosette. The number of NPCs in all rosettes was quantified in two sections of one organoid. All organoids in each group were counted. hESC (H9) derived organoids_batch 1 (control, n = 148; CHIR and SAG, n = 74), hESC (H9) derived organoids_batch 2 (control, n = 175; CHIR and SAG, n = 83), hiPSC #1 (IMR90) derived organoids (control, n = 146; CHIR and SAG, n = 102), hiPSC #2 (GM25256) derived organoids (control, n = 136; CHIR and SAG, n = 77), hiPSC #3 (GM23338) derived organoids (control, n = 195; CHIR and SAG, n = 69). Significance was calculated using an unpaired *t*-test. Center line, median; whiskers, min to max (show all points). (top right) Quantification of VZ thickness by measuring the thickness of VZ from two independent regions in each rosette (at 90-degree angle intervals). The VZ thickness of all rosettes was quantified in two sections of one organoid. All organoids in each group were counted. hESC (H9) derived organoids_batch 1 (control, n = 296; CHIR and SAG, n = 148), hESC (H9) derived organoids_batch 2 (control, n = 350; CHIR and SAG, n = 166), hiPSC #1 (IMR90) derived organoids (control, n = 292; CHIR and SAG, n = 204), hiPSC #2 (GM25256) derived organoids (control, n = 272; CHIR and SAG, n = 154), hiPSC #3 (GM23338) derived organoids (control, n = 390; CHIR and SAG, n = 138). Significance was calculated using an unpaired *t*-test. Center line, median; whiskers, min to max (show all points). (bottom left) Quantification of the rosette size by measuring the diameter of SOX2+ rosette. All rosettes were quantified in two sections of one organoid. All organoids in each group were counted. hESC (H9) derived organoids_batch 1 (control, n =148; CHIR and SAG, n = 74), hESC (H9) derived organoids_batch 2 (control, n = 175; CHIR and SAG, n = 83), hiPSC #1 (IMR90) derived organoids (control, n = 146; CHIR and SAG, n = 102), hiPSC #2 (GM25256) derived organoids (control, n = 136; CHIR and SAG, n = 77), hiPSC #3 (GM23338) derived organoids (control, n = 195; CHIR and SAG, n = 69). Significance was calculated using an unpaired *t*-test. Center line, median; whiskers, min to max (show all points). (bottom right) Quantification of the number of rosettes. The number of rosettes was quantified in three sections of one organoid. All organoids in each group were counted. hESC (H9) derived organoids_batch 1 (control, n =18; CHIR and SAG, n = 15), hESC (H9) derived organoids_batch 2 (control, n = 21; CHIR and SAG, n = 18), hiPSC #1 (IMR90) derived organoids (control, n = 15; CHIR and SAG, n = 18), hiPSC #2 (GM25256) derived organoids (control, n = 15; CHIR and SAG, n = 15), hiPSC #3 (GM23338) derived organoids (control, n = 18; CHIR and SAG, n = 15). Significance was calculated using an unpaired *t*-test. Center line, median; whiskers, min to max (show all points). (d) (left) Representative bright field images of manually-dissected, single rosettes (left; before dissection, right; after dissection). Scale bars, 1 mm. (right) Representative images of single rosettes, immediately after dissection at day 32, immunostained for SOX2 and ZO-1. Magnified images are shown below. Scale bars, 100 μm. (e) Quantification of the size of manually-dissected, single rosettes. Single rosettes were categorized by size based on the length of their diameters. The bar graphs show the relative proportion of the different-sized single rosettes. Normal distribution is represented by the curve; the median diameter is represented by the vertical line. (f) Immunocytochemical analysis of hPSC-derived, reelin-expressing neurons. Scale bars, 100 μm. Quantification of the proportion of VGLUT1+, P73+, and VGLUT+/P73+ neurons are shown on the left (Three biological replicates were evaluated; n = 3). Relative gene expression of *RELN* is shown on the right (Three biological replicates were evaluated; n = 3). Significance was calculated using an unpaired *t*-test. (g) Representative images of single rosettes encapsulated with the RELN+ layer (day 35) immunostained for SOX2 and RELN. Scale bars, 100 μm. (h, i) Immunocytochemical analysis of astrocytes (h) and microglia (i) differentiated from hPSCs. Scale bars, 50 μm for panel ‘h’ and 25 μm for panel ‘i’. (j) Schematic illustration of the glial cell (astrocytes and microglia) microinjection process. Glial cells were injected into intermediate assembloids (day 50), six times per sample, as shown in the image. (k) Representative images of intermediate assembloids (day 50) immediately after being microinjected with glial cells. Astrocytes and microglia were labeled with RFP and GFP, respectively. Scale bars, 100 μm.

Initially, embryoid bodies (EBs) were developed from human pluripotent stem cells (PSCs) through suspension culture. These EBs were cultured in Matrigel for 7 days for neuroectoderm lineage specification. On day 14, Matrigel was removed, and the neuroectoderm structures were cultured under shaking conditions for additional 11 days, resulting in the formation of forebrain organoids with multiple individual neuroepithelium-like structures, known as rosettes, consisting of neural progenitor cells (NPCs). From days 25 to 32, the Hedgehog (Hh) and Wnt pathways ^24,25^ were pharmacologically activated to promote the proliferation and expansion of NPCs—characterized by the increase of SOX2-positive cells, the thickening of the ventricular zone (VZ), and the enlargement of each rosette (Fig. 1b,c and Supplementary Fig. 1a).

On day 32, early forebrain organoids with larger rosettes were manually dissected into single-rosette structures to better mimic the single VZ layer of the developing brain (Fig. 1d and Supplementary Fig. 1b). The resulting single-rosette organoids were uniform and cyst-like structures without considerable variation between different lines (Fig. 1d,e and Supplementary Fig. 1b). In all single rosette organoids, single lumens were observed while maintaining the correct apical polarity crucial for proper neuronal development (Fig. 1d). On day 35, following an additional 3 days of culture to stabilize the dissected structure, five single-rosette organoids whose diameters were closest to the median value were selected and reconstituted using hPSC-derived neurons engineered to express RELN (Fig. 1f,g and Supplementary Fig. 1c). These RELN-expressing neurons exhibited characteristics of glutamatergic neurons, positive for MAP2, VGLUT1, and P73, in addition to the exogenous expression of RELN, with high level of homogeneity (Fig. 1f). Based on the marker expression, the engineered RELN-expressing neurons are likely to function as Cajal-Retzius neurons, which, during human brain development, migrate into the marginal zone (later becoming layer 1) at the early stage of corticogenesis ^17,18^. To provide the necessary cues for precise neural guidance conducive to the formation of structured cortical layers, individual single rosette organoids were encapsulated with a thin Matrigel layer infused with RELN-expressing neurons. The resulting structures were then subjected to an additional culture period of 15 days.

On day 50, glial cell integration into the resulting organoids – forebrain assembloids – was achieved via microinjection of hPSCs-derived astrocytes and microglia into the outer cortical layer (Fig. 1h-k and Supplementary Fig. 1d) ^26,27^. This step aimed to incorporate essential glial cells, contributing to structural and functional maturation. Subsequently, these assembloids were cultured for an additional 30 days to further promote maturation, such as neuronal connection refinement, glial cell maturation, and functional network establishment, resulting in the development of highly uniform human forebrain assembloids closely resembling the complexity and organization of the developing human brain with enhanced cellular diversity and minimized batch effects (Fig. 2a).

**Fig. 2.**
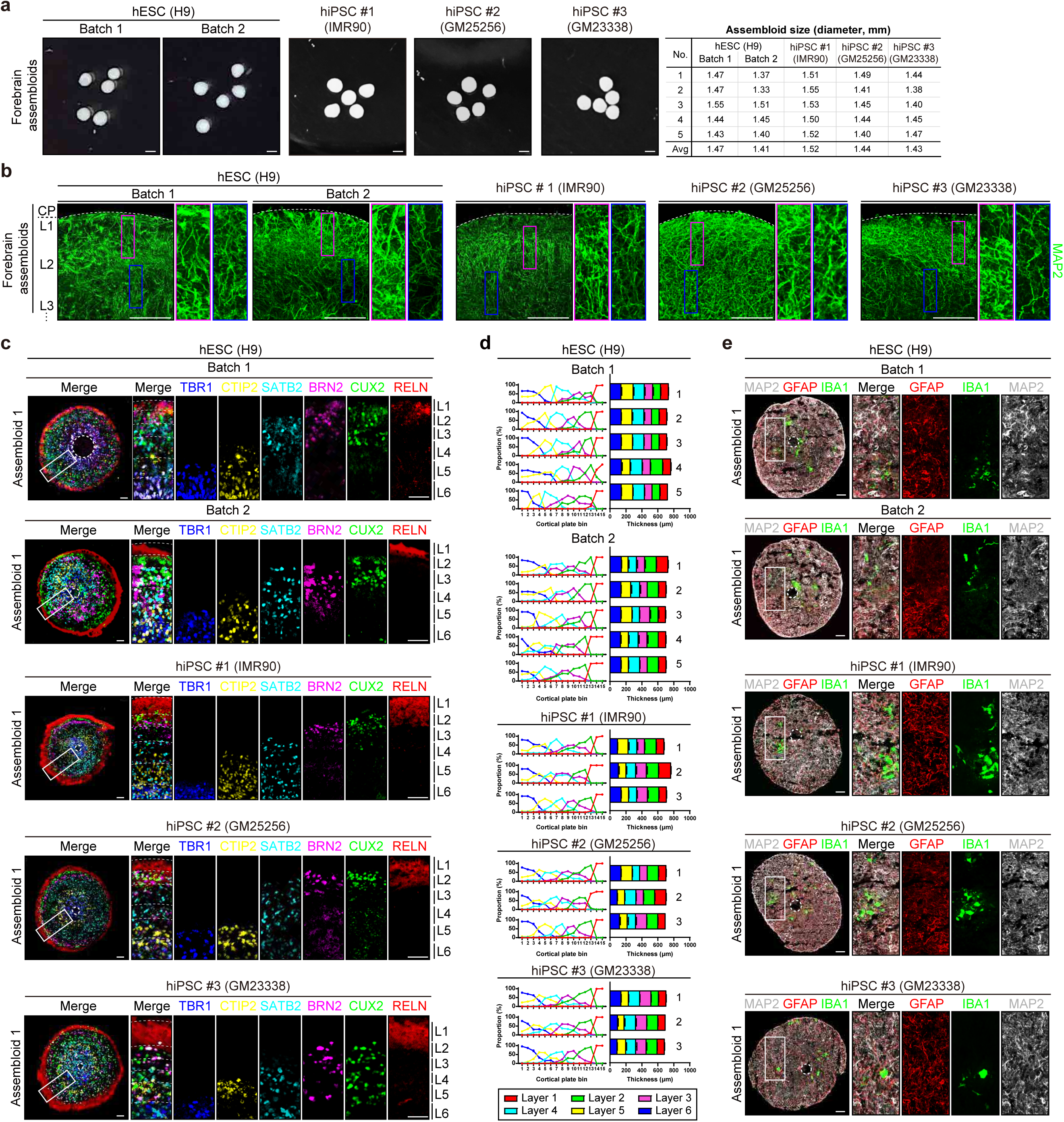
Forebrain assembloids represent the six-layered cortical structure and contain functional glial cells. (a) Representative images of forebrain assembloids (d80). Scale bars, 1 mm. Size measurements of individual forebrain assembloids are shown on the right. The size of the assembloids was determined by averaging the diameters of the assembloids, measured twice at a 90-degree angle. (b) Representative images of forebrain assembloids (d80) immunostained for MAP2 to visualize neuronal projections across cortical layers. Magnified images (insets in the left panels) are shown on the right. CP; cortical plate, L1; layer 1, L2; layer 2, L3; layer 3. Scale bars, 100 μm. (c) Representative images of 6-layered cortical structures of forebrain assembloids (d80). Left panels show merged images of three serial sections at 8-μm intervals in which each section was immunostained for TBR1/CTIP2, SATB2/RELN, and BRN2/CUX2, respectively. Magnified images (insets in the left panels) are shown on the right. Dotted lines demarcate the border of each layer. L1; layer 1, L2; layer 2, L3; layer 3, L4; layer 4, L5; layer 5, L6; layer 6. Scale bars, 100 μm. (d) Quantification of the cortical thickness of each of the 6 layers. The six cortical layers were delineated based on the expression of specific markers associated with each layer of the cortex (TBR1 for layer 6, CTIP2 for layer 5, SATB2 for layer 4, BRN2 for layer 3, CUX2 for layer 2, and RELN for layer 1). For layer quantification, the image of cortical plate of forebrain assembloids is evenly divided into 15 bins, spanning from apical to basal directions. For each bin, the proportion of layer-specific marker-positive cells is calculated as [number of layer-specific marker-positive cells / number of total neurons]. Each bin is assigned to one of the six cortical layers based on the major cell population comprising each bin, determined by the proportion of layer-specific marker-positive cells. The thickness of six cortical layers was measured in four independent regions of each section (at 90-degree angle intervals), and two sections were quantified for one assembloid (n = 8). Five (for hESC derived assembloids) or three (for hiPSC derived assembloids) biological replicates were evaluated. The cortical thickness in individual assembloids is presented as independent bars, labeled with sequential numbers on the graph. (e) Representative images of forebrain assembloids (d80) for neurons (MAP2), astrocytes (GFAP), and microglia (IBA1). Magnified images are shown in the panel on the right. Scale bars, 100 μm.

### Forebrain assembloids mimic the six-layered cortical structure of the human brain

Next, we examined the structural and functional maturity of the forebrain assembloids. The neurons in the newly-developed forebrain assembloids exhibited mature characteristics with pyramidal morphology, as demonstrated by complex dendritic structures and neuronal projections, and displayed spontaneous action potentials and robust neuronal activity at the single-cell level (Supplementary Fig. 2a-d). Moreover, these neurons displayed a radially aligned organization, with axonal projections extending across cortical layers, spanning from the outermost part to the intracortical region of the forebrain assembloids (Fig. 2b and Supplementary Fig. 2a; Supplementary Video 1).

To further evaluate the organization of these neurons into layered structures akin to the human cortex, we conducted immunostaining analysis of forebrain assembloids using layer-specific cortical neuronal markers. We observed in the forebrain assembloids a laminar distribution of layer-specific markers spanning from layer 6 to layer 1 in a proper sequential order (Fig. 2c and Supplementary Fig. 3a,b). Further quantitative analysis of the laminar expression of layer-specific neuronal markers revealed that the newly-developed forebrain assembloids are organized into six stratified cortical layers (Fig. 2d and Supplementary Fig. 3a,b). These cortical layers consist of upper-layer neurons expressing SATB2 (layer 4), BRN2 (layer 3), CUX2 (layer 2), and RELN (layer 1), as well as deep-layer neurons expressing CTIP2 (layer 5) and TBR1 (layer 6). Interestingly, reconstituted neurons expressing RELN consistently remained on the outer surface of the forebrain assembloids, serving as the first neuronal layer, which plays critical roles during the early developmental stages of the human brain (Fig. 2c,d and Supplementary Fig. 3a,b).

In addition to assessing neuronal maturity, we further evaluated the maturity of the forebrain assembloids in terms of glial cell development. We found that glial cells, which were initially localized at the injection site on day 0 (Fig. 1k and Supplementary Fig. 1d), were observed to redistribute throughout the cortex of forebrain assembloids during an extended culture period (Fig. 2e and Supplementary Fig. 4a). GFAP-positive astrocytes displayed distinct star-like, mature branched morphologies and formed tripartite synapses with neurons (Supplementary Fig. 4b). Moreover, IBA1-expressing microglia showed amoeboid to ramified morphologies, indicative of active and resting states, respectively (Supplementary Fig. 4c). These microglia were observed to be in close proximity to neurons and displayed active movement within forebrain assembloids, indicative of functional activity such as the potential for engulfing and remodeling neuronal synapses (Supplementary Fig. 4c and Supplementary Video 2).

Taken together, the structural characteristics of forebrain assembloids suggest a comparable level of structural maturity to the developing human brain at later stages of brain development, wherein radially organized neurons establish the lamination of six cortical layers and glial cells such as astrocytes and microglia are integrated and organized with neurons to form functional networks within the cortex ^18,28,29^.

### Forebrain assembloids represent spontaneous neural activity and functional connectivity

We next investigated the functional maturity of the forebrain assembloids by examining the spontaneous neural activity and functional connectivity. We first assessed synapse formation in forebrain assembloids and observed multiple puncta, marked by co-localization of the presynaptic marker VGLUT1 and the postsynaptic marker PSD95 in the forebrain assembloids, indicating the presence of synapses (Fig. 3a,b and Supplementary Fig. 5a). To further evaluate the spontaneous neural activity of forebrain assembloids, we performed calcium imaging analysis without external stimulation to monitor intracellular calcium dynamics in multiple neurons (Fig. 3c-e and Supplementary Fig. 5b). We observed spontaneous calcium surges in multiple individual neurons, occurring at an average rate of 3-4 surges per minute (Fig. 3c-e and Supplementary Fig. 5b; Supplementary Videos 3,4). In addition to calcium dynamics, we evaluated various electrophysiological properties of forebrain assembloids by performing extracellular recordings using multi-electrode arrays (MEAs). Our analysis demonstrated robust electrical activities in forebrain assembloids, characterized by spontaneous spikes with discrete action potentials and depolarization, as shown in the waveforms (Fig. 3f). Moreover, most of the firing events in forebrain assembloids appeared as bursts, exhibiting repetitive and periodic patterns with an average burst frequency of 0.33 Hz and an interburst interval (IBI) of 3.04 s (Fig. 3g,h and Supplementary Fig. 5c). These data strongly suggest that forebrain assembloids exhibit the robust and spontaneous neural activity.

**Fig. 3.**
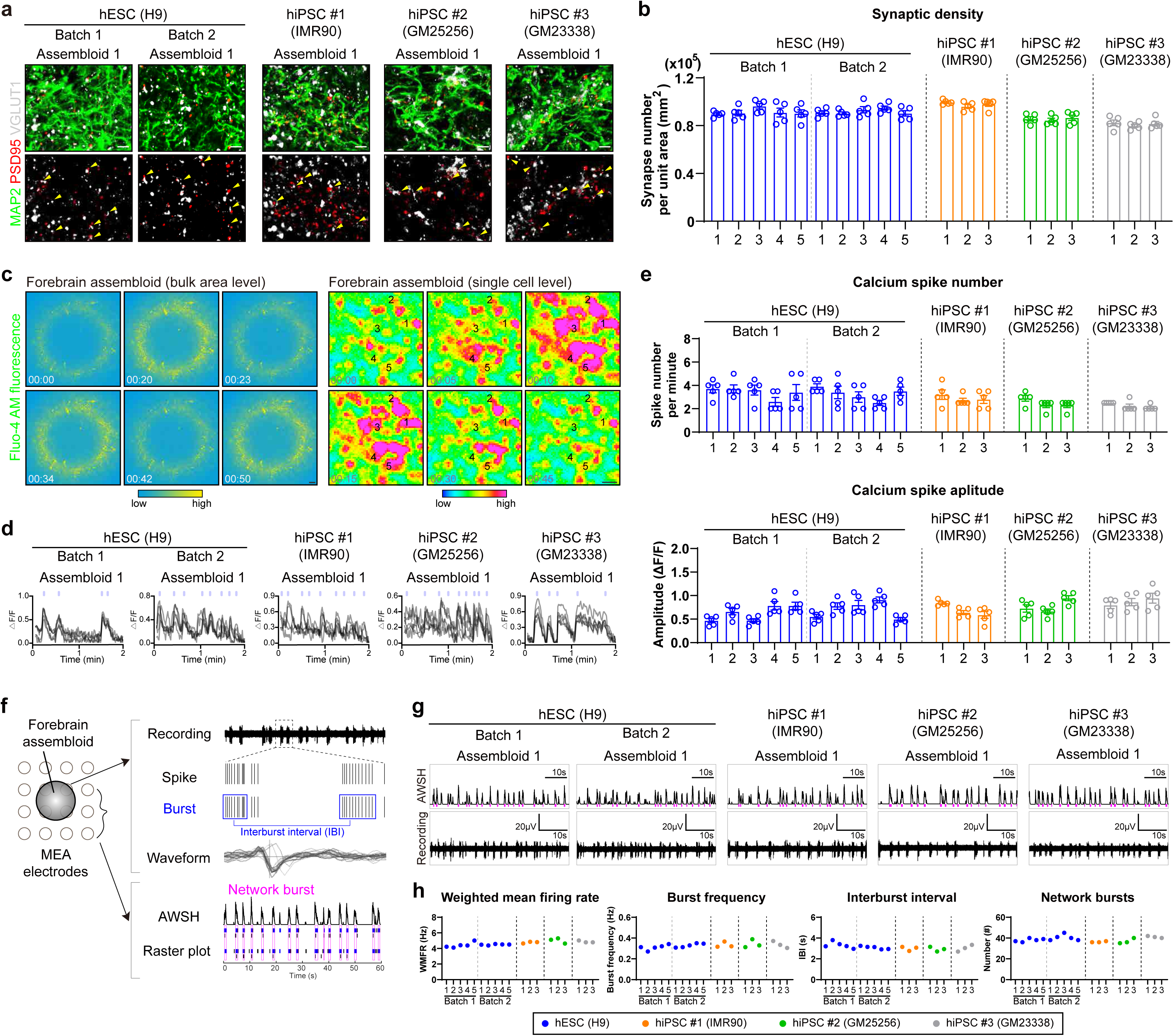
Forebrain assembloids represent the functional connectivity. (a) Representative images of forebrain assembloids (d80) immunostained for neurons (MAP2) and synapses (PSD95 and VGLUT1). The yellow arrowheads indicate synapses co-localized with PSD95 and VGLUT1. Scale bars, 50 μm. (b) Quantification of synaptic density by calculating the number of synapses per unit area (mm^2^) in each section. Five sections were quantified for one assembloid (n = 5). Five (for hESC derived assembloids) or three (for hiPSC derived assembloids) biological replicates were evaluated. The synaptic density in individual assembloids is presented as independent bars, labeled with sequential numbers on the X-axis. (c) (left) Representative images of calcium imaging analysis of forebrain assembloids (d80) at bulk area level. Scale bar, 100 μm. (right) Representative images of calcium imaging analysis of forebrain assembloids (d80) at single cell level. Scale bar, 10 μm. (d) Representative images of calcium imaging analyses of five selected cells in forebrain assembloids. (e) Quantification of spike number per minute (top) and amplitude (bottom) analyzed by calcium imaging of selected cells in forebrain assembloids. The number of spikes and the amplitude were quantified in five selected cells for one assembloid (n = 5). Five (for hESC derived assembloids) or three (for hiPSC derived assembloids) biological replicates were evaluated. The number of spikes and amplitude in individual assembloids is presented as independent bars, labeled with sequential numbers on the X-axis. (f) Schematic representation illustrating the analysis of electrical activity from MEA recordings of forebrain assembloids. Each bar represents a spike. A spike cluster (shown in blue) represents a burst. Burst occurring simultaneously across multiple channels is defined as a network burst (shown in magenta). AWSH, array-wide spike histogram. (g) Representative images of the array-wide spike histogram (AWSH) and recording plot analyzed by MEA in forebrain assembloids. Network bursts are highlighted in magenta. (h) Quantification of the weighted mean firing rate (WMFR), burst frequency, interburst interval (IBI), and the number of network bursts in forebrain assembloids measured by MEA. Recordings were performed for two minutes. The signals from all active electrodes were quantified for each assembloid (n = 1). Five (for hESC derived assembloids) or three (for hiPSC derived assembloids) biological replicates were evaluated. The WMFR, burst frequency, IBI, and the number of network bursts in individual assembloids is presented as independent dots, labeled with sequential numbers on the X-axis.

To investigate whether these active neurons in forebrain assembloids are functionally connected, we analyzed area-wide calcium spike patterns to evaluate the synchrony of calcium surges. We observed highly synchronized calcium activity across the entire area of the forebrain assembloids at a bulk area level (Fig. 3c; Supplementary Video 3). At the single-cell level, individual neurons displayed simultaneous calcium surges within short time windows (1-3s) (Fig. 3c-e; Supplementary Video 4), indicative of functionally connected neuronal networks. This synchronized pattern of neuronal activities was further confirmed through channel-wise MEA analysis. We observed highly synchronized neural activity across multiple channels, displaying network burst events in forebrain assembloids (Fig. 3f-h and Supplementary Fig. 5c). Furthermore, these network bursts showed periodic and regular oscillatory patterns, with an average network burst frequency of 0.3 Hz (Fig. 3h and Supplementary Fig. 5c). These functional characteristics observed in forebrain assembloids, such as synchronized and oscillatory network burst events, mirror the complex spatiotemporal dynamics seen in the developing human brain, wherein synchronous activity of oscillating networks is a prominent feature of functional cortical networks facilitating the coordinated propagation of synaptic signals between functionally related cortical areas ^30–32^.

Taken together, our data from structural and functional analysis demonstrate that the forebrain assembloids represent a mature and interconnected six-layered cortical architecture with cellular diversity, along with the functional synapses and connected neuronal networks.

### Forebrain assembloids recapitulate the cellular composition and transcriptome of the human developing brain at the single-cell level

To examine the maturity and cellular complexity of our newly-developed forebrain assembloids at the single-cell transcriptome level, we performed a comparative analysis of single-cell RNA sequencing (scRNA-seq) data between newly-developed forebrain assembloids and forebrain organoids developed from widely utilized, four independent protocols (Figs. 4,5 and Supplementary Figs. 6,7; Supplementary Data 1). We acquired the scRNA-seq datasets of 13 forebrain assembloids derived from various experimental batches of hESC and 3 different hiPSC lines. Additionally, we obtained publicly available scRNA-seq datasets of four independent, widely used forebrain organoids developed using methods reported in previously published studies ^8,20,33,34^. Single-cell datasets of forebrain assembloids derived from different experimental batches and lines, as well as the datasets from four widely used forebrain organoids were integrated and processed for batch correction. Each dataset was then systematically clustered through principal component analysis (PCA) based on highly variable genes, which led to the identification of distinct clusters. The differential gene expression signatures of the identified clusters in each dataset were then analyzed to assign each cluster to pre-existing, endogenous cell types. This comprehensive approach enabled us to evaluate the maturity, including the cellular complexity, of forebrain assembloids at the level of single-cell transcriptome, in comparison to previously reported organoids.

**Fig. 4.**
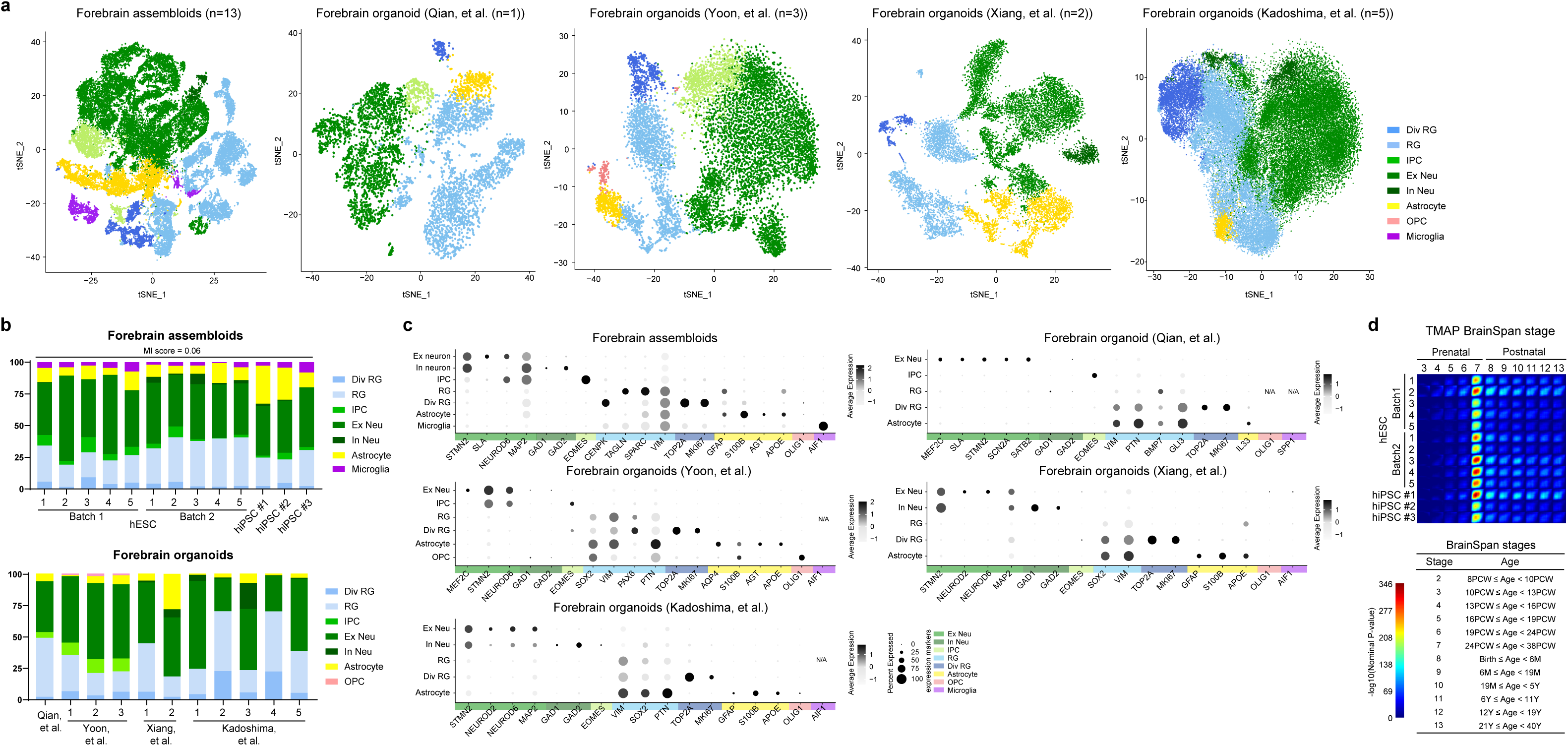
Comparative scRNA sequencing analysis of cell composition and gene transcriptome of forebrain assembloids. (a) tSNE plots of scRNA-seq data from forebrain assembloids (n=13; hESC (H9): five assembloids per batch, two batches analyzed; hiPSC #1 (IMR90), #2 (GM25256), and #3 (GM23338): one assembloid from one batch analyzed) and eleven independent, widely-utilized forebrain organoids, developed using methods by Qian, et al.^20^ (n=1), Yoon, et al.^33^ (n=3), Xiang, et al.^8^ (n=2), and Kadoshima, et al.^34^ (n=5). Cells are colored by cell type and labeled with cell type annotations. RG, radial glia; Div RG, dividing RG; IPC, intermediate progenitor cell; Ex Neu, excitatory neuron; In Neu, inhibitory neuron; OPC, oligodendrocyte progenitor cell. (b) Analysis of the proportion of individual cell types in forebrain assembloids as well as forebrain organoids developed using four different protocols in previous studies. (c) Dot plots showing gene expression of the selected genes expressed in each cluster of current forebrain organoids and forebrain assembloids (circle size, proportion of cells; degree of shading, expression level). (d) Transition mapping (TMAP) of gene expression of forebrain assembloids and human brain tissues from the BrainSpan dataset (compared to stage 2). BrainSpan stages and corresponding ages are shown below. PCW, post conception weeks; M, months; Y, years.

**Fig. 5.**
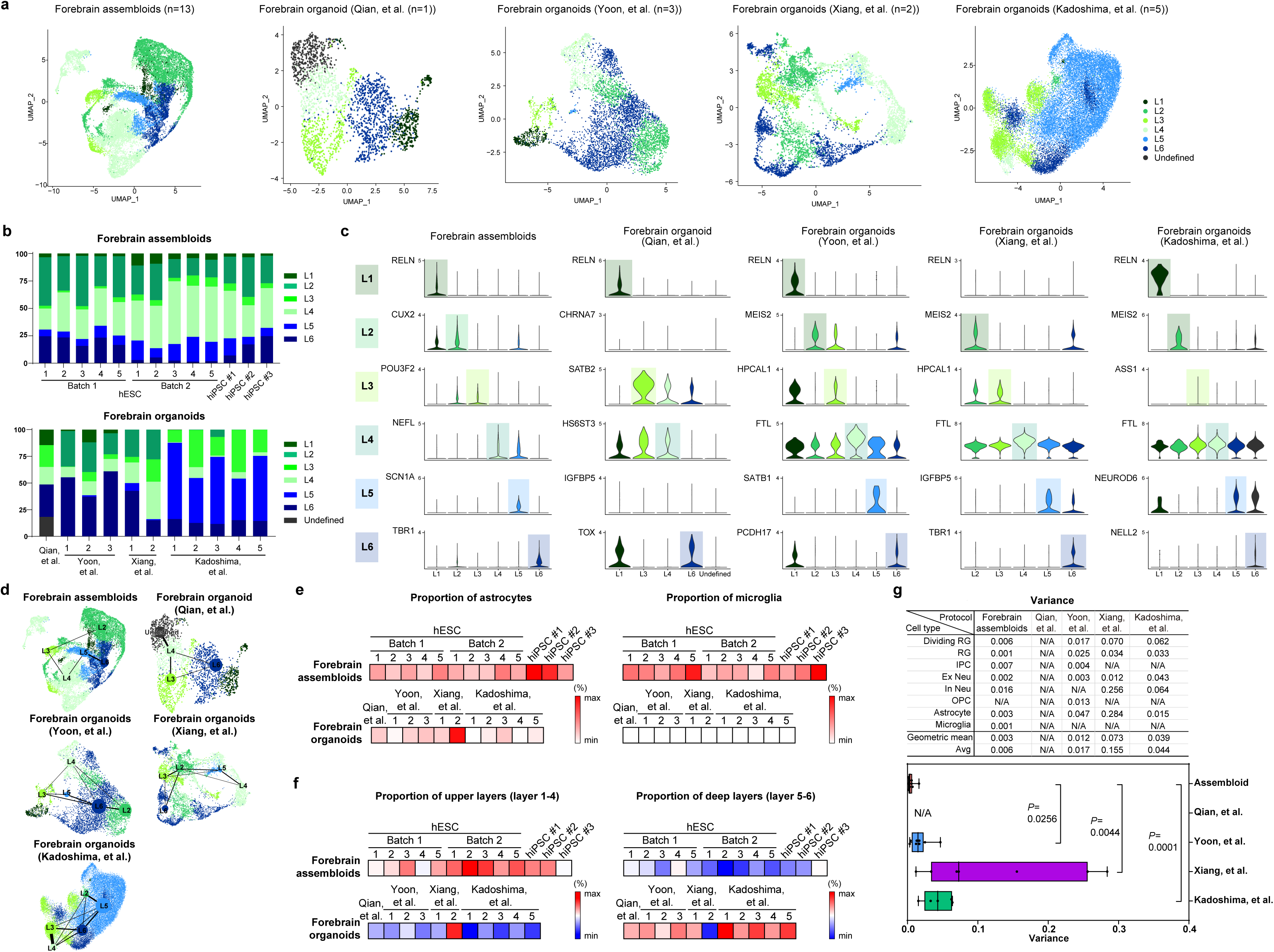
Forebrain assembloids represent the distinct neuronal cell populations of the six cortical layers of the human brain. (a) UMAP plots of scRNA-seq data for clusters marked as excitatory neurons in Fig. 4a, derived from publicly available current forebrain organoids and forebrain assembloids. L1, layer 1; L2, layer 2; L3, layer 3; L4, layer 4; L5, layer 5; L6, layer 6. (b) Analysis of the proportion of individual cells according to the six cortical layers in forebrain assembloids as well as forebrain organoids developed using four different protocols in previous studies. (c) Violin plots showing the selected marker genes expressed in each of the six cortical layers of current forebrain organoids and forebrain assembloids shown in panel ‘a’. (d) Trajectory analysis for excitatory neurons within current forebrain organoids and forebrain assembloids. Cells are colored by six cortical layers. (e) Comparative analysis of the proportion of astrocytes and microglia in forebrain assembloids. The proportions of astrocytes and microglia in forebrain assembloids, in comparison to those in forebrain organoids, are visualized through heatmaps. (j) Comparative analysis of the proportion of upper-layers and deep-layers in forebrain assembloids. The proportions of upper-layers (layer 1-4) and deep-layers (layer 5-6) in forebrain assembloids, in comparison to that of forebrain organoids, are visualized through heatmaps. (g) The variance of each cell type between individual forebrain assembloids, as well as among individual forebrain organoids developed using four different protocols. Center line, median; whiskers, min to max (show all points).

Our analysis showed that all forebrain assembloids exhibited a high degree of consistency, containing seven transcriptionally distinct cell types within forebrain lineages, including radial glia (RG), dividing RG, intermediate progenitor cells (IPCs), excitatory neurons, inhibitory neurons, astrocytes, and microglia (Fig. 4a-c and Supplementary Fig. 6; Supplementary Data 1). Notably, analysis of cellular compositions and transcriptional profiles of forebrain assembloids demonstrated a high correlation with the human fetal brain at late fetal periods (Fig. 4d). Further in-depth analysis revealed that the neurons in forebrain assembloids were segregated into six distinct cell populations expressing layer-specific markers, with each population correlated to a specific cortical layer of the human brain ^35,36^ (Fig. 5a-c and Supplementary Fig. 7). Moreover, the trajectory analysis demonstrated that the excitatory neurons in each cortical layer of forebrain assembloids followed a normal developmental process, sequentially generating layer-specific neurons from layer 6 to layer 2 (Fig. 5d), closely resembling the developmental pattern observed in the human cortex during development ^37^. This finding strongly suggests that forebrain assembloids are comparable to the human developing brain in terms of cortical layer specification and cellular compositions.

In comparison to forebrain assembloids, analysis of scRNA-seq datasets from four independent, widely used forebrain organoids reported in previous studies ^8,20,33,34^ revealed that, although the cell types and their proportions varied across protocols, these forebrain organoids were primarily composed of RG, dividing RG, and excitatory neurons, with a small proportion of IPCs as well as few other cell types (Fig. 4a-c and Supplementary Fig. 6; Supplementary Data 1). Specifically, forebrain organoids developed using methods by Kadoshima, et al.^34^ or Xiang, et al.^8^ exhibited a high level of heterogeneity between batches, and predominantly consisted of excitatory neurons, RG, dividing RG, with a minor population or absence of inhibitory neurons and astrocytes (Fig. 4a-c). The majority of neurons in these organoids showed characteristics typical of deep-layer neurons, comprising layer 5 and layer 6 neurons, with relatively fewer upper-layer neurons (layers 2, 3, and 4), and the absence of layer 1 neurons (Fig. 5a-c and Supplementary Fig. 7). Similar deep-layer properties were also observed in forebrain organoids developed from Yoon, et al.^33^, exhibiting a high proportion of early-born layer 6 neurons with a relatively lower proportions of upper-layer neurons (Fig. 5a-c and Supplementary Fig. 7). Excitatory neurons in forebrain organoids developed from Qian, et al.^20^ also exhibited deep-layer properties with 30% of layer 6 neurons, alongside smaller proportions of upper-layer neurons, including 15% of layer 1, 21% of layer 3, and 16% of layer 4 neurons, with the absence of layer 2 and 6 neurons (Fig. 5a-c and Supplementary Fig. 7). Furthermore, a minor proportion or absence of astrocytes was observed in all forebrain organoids, except for one batch of the organoids developed from Xiang, et al.^8^ which exhibited a high degree of heterogeneity between batches, with the complete absence of microglia. Notably, in contrast to forebrain assembloids, the pseudotime trajectory analysis revealed that the excitatory neurons in each cortical layer of forebrain organoids followed a developmental process that did not correspond to the sequential development of cortical layers observed during the human cortex development (Fig. 5d). Taken together, these data suggest that widely-used forebrain organoids reported in previous studies consist of a high proportion of deep-layer neurons with relatively lower populations of late-born neurons, alongside a minor population or absence of glial cells such as astrocytes and microglia.

Comparative analysis was further performed to assess the level of maturity of forebrain assembloids compared to previously reported forebrain organoids. We found that our forebrain assembloids contained a greater proportion of glial cells, including astrocytes and microglia. Specifically, astrocytes and microglia constituted approximately 13% and 4% of the total cell population, respectively, rendering forebrain assembloids more comparable to the human developing brain than previously reported forebrain organoids with lower proportions of astrocytes, as well as the complete absence of microglia (Fig. 5e). These results strongly suggest that forebrain assembloids exhibit enhanced maturity at the level of cellular compositions.

In addition, forebrain assembloids exhibited all six distinct cortical layers with a reduced presence of deep layer neurons (layers 5 and 6) and a significantly higher proportion of upper layer neurons (layers 1, 2, 3, and 4) (Fig. 5f). Although their proportions varied across protocols, excitatory neurons in previously reported forebrain organoids were predominantly composed of a deep-layer neurons (layers 5 and 6) with relatively lower levels of upper-layer neurons, alongside the absence of certain layer-specific upper-layer neurons, leading to an incomplete alignment with all six cortical layers (Fig. 5b,f).

Lastly, mutual information (MI) score^38^ was calculated across samples to evaluate the variability of our forebrain assembloids derived from multiple batches and different lines, revealing that all forebrain assembloids demonstrated a high level of consistency (Fig. 4b). In contrast, currently available forebrain organoids developed using the same protocols exhibited a comparatively higher degree of variability between batches and individual organoids within the same methodology, as demonstrated by the variance test (Fig. 5g). These data suggest high reproducibility and low variation of forebrain assembloids across batches and lines, further implying a high level of consistency and uniformity without significant batch effects and heterogeneity.

Taken together, these results demonstrate that forebrain assembloids comprise the six distinct cortical layers of neurons and glial cells with a high degree of consistency, suggesting a level of maturity and cellular complexity highly comparable to that of the cortex in the developing human brain (Fig. 5 and Supplementary Figs. 6,7).

## Discussion

A major conceptual advance presented in our current work is the development of effective strategies to generate newly conceived reconstituted brain organoids called “brain assembloids”, which are defined as “organoids created by reconstituting multiple cell types of human tissues (not an alternative definition as ‘hybrid organoids’)”, with multiple cell types that structurally and functionally recapitulate the pathophysiology of human brains. To this end, two critical signaling molecules involved in embryonic patterning, Hedgehog and Wnt, were employed to increase neural progenitor populations at an early stage of organoid formation, based on previous findings that these two patterning cues induce cellular proliferation of neuronal progenitor cells, leading to the formation of embryonal tumors with multilayered rosettes (ETMR) ^24,25^. The mature patterning of the forebrain cortex, leading to the thickened laminar organization of cortical layers, was achieved by subsequently reconstituting early single-rosette brain organoids with a first layer of neuronal cells, called Cajal–Retzius cells, which were engineered to express RELN after being differentiated from hPSCs. Conventional brain organoids lack this important first layer at the early stage of development, during which RELN plays a critical role as a patterning cue for neural guidance, in which newly-born neurons pass the early-generated layers and migrate beneath the RELN-expressing outer layer to generate an ‘inside-out’ six-layered structure with laminar organization of cortical layers ^39^. By integrating the RELN-expressing neuronal layer into organoids with an increased population of NPCs induced by Hh and Wnt, we successfully achieved mature laminar organization of forebrain cortical layers. At the later stage, the resulting organoids were further microinjected with glial cells including astrocytes and microglia to generate the final form of forebrain assembloids, which represent dynamic cellular interplay between neurons and glial cells, and thus provide a powerful tool to understand new mechanistic insights into how particular signaling feedback between multiple cell types in the human brain is regulated under normal and pathological conditions.

It is important to note that one of the conceptual advances in our study is generating consistent and uniform, cyst-like structure of forebrain assembloids with a single ventricular zone and 6 cortical layers. We have achieved this uniformity without considerable variation between organoids and different batches by manually dissecting brain organoids of the early stage to generate single rosettes whose sizes were relatively bigger and consistent due to the treatment with Hh and Wnt agonists. This step was followed by the reconstitution procedure of a uniform single rosette, which contains a single lumen that becomes the ventricular zone later, with the first layer of RELN-expressing neurons and glial cells, where high levels of experimental control were also achievable. In this way, we were able to create highly consistent and uniform assembloids without significant batch effects and to solve the major heterogeneity issues which are mostly due to the difficulty in controlling the number and the size of each rosette in current multi-rosette organoids.

Two recent papers have described single rosette-based protocols for developing brain organoids ^40,41^. These previous works utilized two-dimensional (2D), structurally premature rosettes developed through 2D neuroepithelial culture. A conceptual advance of our study over these earlier works lies in utilizing 3D, uniform rosettes developed by 3D brain organoid culture at the early stage, which yields not only uniform but also highly reproducible assembloids, eliminating batch effects and heterogeneity. More importantly, we developed consistent and uniform, cyst-like structure of forebrain “assembloids” by employing a modular and stepwise cellular reconstituting method, incorporating engineered RELN-expressing neurons and glial cells. This unique approach induced the development of a mature cortex structure and cellular diversity in our forebrain assembloids, which was not achieved in the previous studies.

Taken together, our study, which has the potential to profoundly affect our understanding of the role of complex cellular interactions in the development of various brain disorders, will facilitate the establishment of an innovative model platform to study a range of human neurological diseases, whose understanding of pathogenesis requires an organoid system that is capable of representing mature characteristics of functional human brains, including dynamic interplay between neurons and other cell types and will further provide a unique tool for the development of new therapeutic options that can be customized for individual patients.

## Methods

### hPSC culture

hESCs (H9) and hiPSCs (IMR90, female) were obtained from WiCell. hiPSCs (GM25256, male; GM23338, male) were obtained from the National Institute of General Medical Sciences Cell Repository through the Coriell Institute for Medical Research. All hPSCs were maintained on mitomycin C-treated mouse embryonic fibroblasts (MEFs) in hPSC medium, containing DMEM/F12 (Gibco) supplemented with 20% KnockOut Serum Replacement (Gibco), 1× Glutamax (Gibco), 1× Non-essential amino acids (Gibco), 1% penicillin–streptomycin, 100 μM 2-Mercaptoethanol (Sigma) and 10 ng/ml human bFGF (Peprotech). All hPSC lines were cultured on each well of a 24-well plate with 0.5 ml of culture media (with 5 x 10^4^ feeder cells per well). The cells were fed daily and passaged by manual dissection at 70% confluence (10 manually-dissected, small colonies of hPSCs were plated on each well of a 24-well plate). Cells used in this study were negative for mycoplasma contamination (e-Myco Mycoplasma PCR detection kit).

### *In vitro* differentiation of neurons, astrocytes and microglia

Glutamatergic neurons were differentiated from hPSCs using the protocol modified from the previous work ^42^. In brief, hPSC colonies were detached from the feeder layer using collagenase IV (Thermo) at 37 ℃ for 1 h. 10-16 PSC colonies (5-8 hPSC colonies from each well of 24-well plate, 2 wells) were collected in 0.5ml of N2/B27 media, consisting of DMEM/F12, 1× N2 supplement (Gibco), 1× B27-RA supplement (Gibco) and 1× penicillin–streptomycin, and plated on 35 mm petri dish. On day 1, the medium was changed with N2/B27 medium supplemented with 10 μM SB431542 (Sigma) and 0.1 μM LDN193189 (Stemgent). On day 7, EBs were transferred to Matrigel-coated (10% Matrigel in DMEM/F12, Growth Factor Reduced; Corning) plates and cultured in N2/B27 medium with 10 nM SB431542. On day 16, cells with rosette structures were collected and dissociated mechanically by pipetting. Dissociated cells were plated on Matrigel-coated plates in NPC medium, consisting of DMEM/F12, 1× N2 supplement, 1× B27-RA supplement, 20 ng/ml basic FGF and 1 μg/ml laminin (Thermo). At 80% confluency, NPC medium was changed into neuronal medium, consisting of DMEM/F12, 1× N2 supplement, 1× B27-RA supplement, 1× penicillin–streptomycin, 20 ng/ml BDNF (Peprotech), 10 ng/ml GDNF (Peprotech), 250 μg/ml dibutyryl cyclic-AMP (Biogems) and 200 nM L-ascorbic acid (Sigma). The cultures were maintained in neuronal medium for 2 weeks on 10 μg/ml poly-L-ornithine (Sigma) and 5 μg/ml laminin-coated plates. Neuronal identity was validated by qRT-PCR and immunocytochemistry.

hPSC-derived astrocytes were generated as previously described ^27^. Briefly, hPSC-derived NPCs were differentiated into astrocytes by seeding dissociated single cells at 15,000 cells/cm^2^ on Matrigel-coated plates in Astrocyte medium (ScienCell: 2% FBS, Astrocyte Growth Supplement, 1% penicillin–streptomycin in astrocyte basal medium) and maintained. After 30– 40 d of differentiation, hPSC-derived astrocytes were characterized using qRT-PCR and immunocytochemistry.

hPSC-derived microglia were generated as previously described ^26^. Briefly, hPSC colonies were detached from the feeder layer with collagenase IV for 1 h. hPSC colonies were collected in microglia medium consisting of 10 ng/ml IL-34 (Peprotech) and 10 ng/ml GM-CSF (Peprotech) in 100-mm sterile petri dishes. After 7–14 d of differentiation, EBs with a cystic morphology were selected and transferred to 10 μg/ml poly-L-ornithine and 5 μg/ml laminin-coated plates. Six additional titrations were performed every 5 d. Further maintenance was performed in microglia medium with 100 ng/ml IL-34 and 5 ng/ml GM-CSF. Microglial identity was validated by qRT-PCR and immunocytochemistry.

### Generation of RELN-expressing neurons

The lentiviral construct for RELN expression was generated by subcloning domains 3–6 (central domains) of *RELN* from the pCrl construct (Addgene #122443, kindly gifted by Mi-Ryoung Song, Gwangju Institute of Science and Technology) into the pLenti 6.3-DEST (Thermo) lentiviral expression vector. NPCs were transduced with the *RELN*-containing lentivirus using protamine sulfate (5 μg/ml). Three days after transduction, NPCs were expanded in NPC medium, containing blasticidin (6 μg/ml), for antibiotic selection. NPCs were then differentiated into glutamatergic neurons. qRT-PCR was performed to confirm RELN expression.

### A stepwise development of forebrain assembloids

hPSCs were cultured on a feeder layer in a 24-well plate (5-8 hPSC colonies in each well of the 24-well plate). When the diameter of hPSC colonies reached 1.0–1.5 mm, the hPSC medium was replaced with 0.25 ml of 1 mg/ml collagenase IV and incubated at 37 °C for 1-2 h to detach the colonies from the feeder layer without disrupting their colony structure. Detached colonies were transferred to a 15-ml tube and washed with 1 ml of hPSC medium. 2 ml of EB medium, comprising hPSC medium (without bFGF) supplemented with 2 μM Dorsomorphin (Sigma) and 2 μM A83-01 (Tocris), was added, and the 5-8 hPSC colonies (from each well of the 24-well plate) in EB medium were transferred and cultured in a 35-mm petri dish at 37 ℃ to form EBs. On day 5, half of the medium was replaced with neural induction medium consisting of DMEM/F12, supplemented with 1× N2 supplement, 10 μg/ml Heparin (Sigma), 1× penicillin–streptomycin, 1× Non-essential amino acids, 1× Glutamax, 1 μM CHIR99021 (Tocris), and 1 μM SB-431542 (Cellagentech). On day 7, 5-8 EBs were transferred to a 1.5 ml microcentrifuge tube and mixed with 50 μl of Matrigel and 33 μl of neural induction medium. A total of 88 μl of mixture containing EBs was plated onto the center of a 35-mm petri dish and incubated at 37 ℃ for 30 min. Then, 2 ml of neural induction medium was added, and EBs were cultured until day 14 to induce neuroepithelium-like structures. On day 14, the Matrigel-embedded neuroepithelium structures were mechanically dissociated by gentle pipetting, and the resulting structures were then transferred to a new 35-mm petri dish containing 2 ml of differentiation medium consisting of DMEM/F12 supplemented with 1× N2 supplement, 1× B27 supplement, 1× penicillin–streptomycin, 1× 2-Mercaptoethanol, 1× Non-essential amino acids, 1× Glutamax, and 2.5 μg/ml Insulin (Sigma). These neuroepithelium structures were cultured in differentiation medium until day 25 in a shaking incubator (Eppendorf; New Brunswick S41i, 90 rpm) to form forebrain organoids with multiple rosettes.

From days 25 to 32, the organoids were cultured in differentiation medium supplemented with 1 μM CHIR99021 and 400 nM SAG (Millipore). Rosette structures in 32-day-old forebrain organoids were observed under an inverted microscope at 10X magnification (Evos XL Core, Invitrogen), and manually dissected using fine forceps (Fine Science Tools). About 5-7 single-rosette organoids (about 25-40 single-rosette organoids derived from SCZ patients) were generated from one forebrain organoid. Since each batch of hPSCs typically yields approximately 5-8 forebrain organoids, this results in the generation of approximately 25-56 single-rosette organoids per batch (around 125-320 single-rosette organoids derived from SCZ patients). After dissection, the single-rosette organoids were cultured for an additional 3 days to stabilize the dissected structures.

On day 35, five single-rosette organoids, whose diameters were closest to the median values within the range of 25-40 single-rosette organoids, were selected for further procedures. For hESC-derived normal assembloids, two batches were used in each set of experiments (a total of 10 single-rosette organoids were generated and 10 final assembloids were analyzed in each experiment). For hiPSC-derived normal assembloids, one batch was used (a total of 5 single-rosette organoids were generated, and 3 final assembloids were analyzed in each experiment). For SCZ-derived patient assembloids, one batch was used (a total of 5 single-rosette organoids were generated, and 5 final assembloids were analyzed in each experiment). The resulting single-rosette organoids were then encapsulated with 1–2 μl of Matrigel containing RELN-expressing neurons at a concentration of 1 × 10^4^ cells/μl and cultured in 2 ml of differentiation medium until day 50.

On day 50, hPSC-derived astrocytes (1 × 10^4^ cells) and microglia (2 × 10^3^ cells) ^35^ were then microinjected into the outer cortical layer of 50-day-old forebrain assembloids. Cells were microinjected at six different locations, which are evenly distributed through the outer cortical layer, with 50 μm inside the surface of the cortical layer. The resulting assembloids were cultured for an additional 30 days in 2 ml of maturation medium consisting of Neurobasal medium (Gibco) supplemented with 1× B27 supplement, 1× penicillin–streptomycin, 1× 2-Mercaptoenthanol, 0.2 mM Ascorbic Acid, 20 ng/ml BDNF (Peprotech), 20 ng/ml GDNF (Peprotech), and 0.5 mM cAMP (Sigma) to generate the final form of forebrain assembloids. From days 14 to 80, all cultures were maintained in a shaking incubator with medium changes every other day.

### Lentivirus production

Lentivirus production was performed as previously described ^23^. In brief, transfection mixtures were prepared by mixing 9 μg of packaging vector (gag, pol; pCMV.dR 8.74), 3 μg of envelope vector (VSV-G; pMD2.G), 10 μg of the transfer vector of interest and three volumes of TransIT-LT1 Transfection Reagent (Mirus). 48 h after transfection, lentivirus-containing supernatants were collected and filtered through a 0.45-μm filter. The virus-containing supernatants were further concentrated by centrifuging at 24,000 rpm for 2 h at 4 °C.

### Lentiviral infection in forebrain assembloids

Forebrain assembloids were incubated in assembloid medium (2 ml) containing EGFP lentivirus (1.0 x 10^6^ TU/ml, 200 μl) with polybrene (10 μg/ml) for 3 h at 37 °C. The virus-containing medium was removed, and assembloids were washed with warm DPBS twice. Assembloids were cultured for 3 days more in assembloid medium and analyzed by fluorescence imaging.

### qRT-PCR

Total RNA was extracted from multiple cells and organoids as previously described ^23^. In brief, cells and organoids were homogenized by trituration and trypsinization. RNA was extracted using the RNeasy Plus Mini Kit (QIAGEN) and first-strand cDNA was synthesised using a High-Capacity cDNA Reverse Transcriptase Kit (Applied Biosystems) with oligo dT. qRT-PCR was performed using SYBR Green Supermix (Applied Biosystems) and a One-step Cycler (Applied Biosystems). Gene expression was normalized to the housekeeping gene GAPDH.

### Immunohistochemistry

Immunohistochemistry was performed as previously described ^23^. In brief, samples were fixed in 4 % paraformaldehyde (PFA) for 15 min and cryopreserved in 30 % sucrose overnight. Samples were embedded in an OCT compound (Sakura) and frozen at −20 ℃. Then, 8–20-μm-thick sections were generated using a cryostat (Leica). Frozen sections were fixed in 4 % PFA for 20 min at 4 ℃, washed with PBS three times and blocked in 2 % goat serum and PBS containing 0.25 % Triton X-100 (PBS-T) for 1 h at RT. The sections were then incubated with primary antibodies diluted in blocking buffer overnight at 4 ℃. The following primary antibodies were used: TUJ1 (1:300, BioLegend), SOX2 (1:300, Abcam), CTIP2 (1:300, Abcam), CUX2 (1:300, Abcam), SATB2 (1:300, Abcam), TBR1 (1:300, Abcam), MAP2 (1:300, Abcam), GFAP (1:300, Dako), IBAI (1:60, Santacruz), RELN (1:200, MBL), BRN2 (1:300, Santacruz), PSD95 (1:300, Invitrogen), VGLUT1 (1:100, Santacruz), ZO-1 (1:200, Santacruz), and P73 (1:200, Thermo). Sections were washed three times with 0.25 % PBS-T and incubated with secondary antibodies (1:1,000, Life Technologies) diluted in blocking buffer for 1 h at RT. The sections were washed with 0.25 % PBS-T and mounted with Prolong Gold mounting reagent (Invitrogen).

For immunocytochemistry, cells were plated on 10 μg/ml poly-L-ornithine and 5 μg/ml laminin-coated coverslips on a 12-well plate. When reached 80% confluence, cells were washed with PBS and fixed in 4 % PFA for 5 min at RT. Cells were washed with PBS three times and blocked for 40 min at RT. Cells were then incubated with diluted primary antibodies for 1 h at RT and washed three times with PBS-T. Cells were then incubated with secondary antibodies diluted in blocking buffer for 40 min at RT. The cells were washed two times with PBS-T and mounted on glass slides.

### Multi-electrode array (MEA) recording

24-well MEA plates (Axion Biosystems, Atlanta, GA, USA) were coated with 10 μg/ml poly-L-ornithine and 5 μg/ml laminin solution prior to assembloid culture. Assembloids were placed on MEA plates and cultured for 2 weeks to be attached to the electrodes with media change in every 3 days. After two weeks of attachment and stabilization, recordings were captured to measure basic parameters such as weighted mean firing rate, burst frequency, mean interburst interval, and the number of network bursts using a Maestro MEA system and AxIS Software Spontaneous Neural Configuration (Axion Biosystems). Spike detection was carried out using the AxIS software by setting an adaptive threshold at 5.5 times the noise’s standard deviation estimated for each channel (electrode). Before recording, the plate was left to stabilize for 10 minutes inside the Maestro device.

The electrodes with at least 5 spikes/min were defined as active electrodes. Bursts within the data from each electrode were recognized based on an inter-spike interval (ISI) threshold requiring a minimum number of 5 spikes with a maximum ISI of 100 ms. A minimum of 12 spikes under the same ISI criterion with a minimum of 50 % active electrodes were required for network bursts in the well. Raster plot and array-wide spike histogram were obtained using Axion Biosystems’ Neural Metrics Tool.

### Whole-cell patch-clamp recording

Whole-cell patch-clamp recording was performed from sections sectioned from forebrain assembloids. AAV1-hSyn1-GFP (Addgene #50465) was diluted in DPBS on ice and microinjected five times into the forebrain assembloids. Assembloids were embedded in 4% low melting point agarose and sliced into 200 μm-thick sections on a vibratome. Slices were recovered in oxygenated aCSF containing 119 mM NaCl, 2.5 mM KCl, 2.5 mM CaCl_2_, 2 mM MgSO_4_, 1.25 mM NaH_2_PO_4_, 26 mM NaHCO_3_ and 10 mM D-glucose for 1 h, equilibrated with 95 % O_2_ and 5 % CO_2_ (pH 7.3 ~ 7.4) at RT. After recovery, slices were transferred into a recording chamber perfused with aCSF solution. Whole-cell post-synaptic patch-clamp recordings were performed using a MultiClamp 700B amplifier (Molecular Devices). Recording glass pipettes (4-9MΩ) were filled with an internal solution containing 120 mM K-Gluconate, 5 mM NaCl, 0.2 mM EGTA, 1 mM MgCl2, 10 mM HEPES, 2 mM MgATP, and 0.2 mM NaGTP (adjusted to pH 7.2 with KOH). All the recorded cells showed GFP expression. Membrane properties and excitability were measured and analyzed with Clampfit 10.1 software (Molecular Devices).

### Calcium imaging

Forebrain assembloids were incubated with organoid medium containing 1 μM Fluo4-AM (Invitrogen) for 3 h at 37 ℃. Assembloids were washed once with DPBS and incubated in brain organoid medium. Time-lapse image sequences were acquired for 2 min with 1 sec intervals on a Nikon confocal microscope. ΔF/F traces in the selected cells were calculated and shown in the graph (ΔF/F=(F – F0)/F, where F is the fluorescence at given time point and F0 is the minimum fluorescence of each cell). The average amplitudes (ΔF/F) and frequencies of spikes detected in the forebrain assembloids are 0.6-1.4 and 3-4 spikes/min. Spontaneous calcium activities were analyzed with ImageJ software.

### Single-cell RNA sequencing

#### Cell harvest

Forebrain assembloids were collected and washed twice with DPBS. The cells were treated with Accutase for 2–3 minutes at RT before being dissociated into single cells by pipetting and centrifugation at 300g for 5 minutes. The cells were resuspended in the medium (DMEM with 10% FBS) and strained through a 40-μm cell strainer. Each sample was run on 10× Chromium Single-Cell Chips (10X GENOMICS) following the manufacturer’s instructions.

#### Single-cell RNA library preparation and sequencing

The scRNA-seq library was prepared using the Chromium Single Cell 3 Prime platform (v3.1 Chemistry;10x Genomics) according to the manufacturer’s instructions. Briefly, 10,000 cells per sample were loaded into the Chromium Controller in order to generate single Gel Bead-in-Emulsions (GEMs) with the Chromium Next GEM Single Cell 3 Prime Reagent Kit v3.1 (PN-1000268, 10x Genomics). The cells were lysed and released RNAs were synthesized into cDNA through reverse transcription in individual GEMs. Full-length cDNA was synthesized by capturing polyadenylated mRNA with poly(dT) primers and barcoding it. cDNA was synthesized by incubating at 53°C for 45 minutes and 85°C for 5 minutes. cDNA amplification was performed for 12 PCR cycles following GEM cleanup to generate sufficient amounts of DNA for library construction. Single Cell 3’ GEX and feature barcode libraries were sequenced on the Illumina NovaSeq 6000.

#### Pre-processing of Single-Cell RNA-sequencing data

Cell Ranger pipeline (v6.1.2 10X Genomics) ^43^ was used to demultiplex samples, process barcodes, and align reads to the GRCh38 reference genome provided by Cell Ranger (v6.1.2). The ‘mkfastq’ module of Cellranger was used to demultiplex raw base call (BCL) data produced by Illumina sequencers into FASTQ files. The ‘count’ module of Cellranger was used for alignment, filtering, barcode and UMI counting. Single-cell gene count matrices were constructed for each sample using Seurat (v4.3.0) ^44^ with cells that had number of detected features >= 500 and features that had number of detected cells >=5. Given varying distribution of nCount_RNA, nFeature_RNA and percent.mt among respective sample, we implemented distinct maximum cutoff thresholds (nCount_RNA <50,000~300,000, nFeature_RNA < 9,000~12,000 and percent.mt < 5 ~ 15). To remove cells considered as doublets, scDblFinder (v1.14.0) ^45^ detected doublets using the expected doublet rate of 0.8%.

Single-cell gene count matrices were merged and log-normalized using ‘NormalizeData’ function. 2000 top highly variable genes were selected based on average expression and dispersion for each gene with vst selection methods using the ‘FindVariableFeatures’ function. All respective samples were feature-level scaled using ‘ScaleData’ to reduce the effects of outliers.

To remove batches derived from each sample, ‘RunFastMNN’ was used to integrate Seurat object using batchelor (v1.13.3) ^46^. When excitatory neurons were extracted from the object and pre-processed, the anchors between 13 datasets were identified using ‘FindIntegrationAnchors’ and utilized to integrate all datasets through ‘IntegrateData’.

For the analysis of scRNA-seq data of forebrain organoids, publicly available single-cell datasets from eleven independent, widely utilized forebrain organoids, developed using four different methods in previous studies (Qian, et al.^20^, n=1; Xiang, et al.^8^, n=2; Yoon, et al.^33^, n=3; Kadoshima, et al.^34^, n=5) were obtained from GEO database (GSE137941; GSE98201, GSE107771, and GSE132672, respectively). For forebrain organoids developed using methods by Qian, et al., Xiang, et al., or Yoon, et al., single-cell datasets generated by the original publications were used. For organoids developed using the method by Kadoshima, et al., single-cell datasets generated by Bhaduri, et al.^47^ were used. Seurat objects for these datasets were created using features detected in at least 10 cells and cells having at least 500 features. Due to the diverse distribution of percent.mt across different samples, we applied specific maximum cutoff thresholds ranging from 5 % to 15 % for each sample. The remaining processes, including the removal of doublets, pre-processing, and batch correction, were conducted under the same conditions as described above.

#### Clustering

The nearest neighbors present in the single cell data were found with ‘FindNeighbors’ function using dimensions from NMF. Graph-based Louvain clustering was done with ‘FindClusters’ function. Thereafter, the 50 NMFs were used for t-distributed Stochastic Neighbor Embedding (t-SNE) non-linear dimensionality reduction using RunTSNE.

#### Cell-type annotation

Cell-type annotations of identified clusters of current forebrain organoids and forebrain assembloids were performed by comparison of marker genes in each cluster to those of previously annotated cell types ^47^. The genes used to annotate each cluster are provided in Supplementary Data 1. When marker-gene based annotation was not available, literature-based annotation was used.

#### Annotation of six cortical layers

Excitatory neuron clusters are re-clustered at a resolution of 1.0 and annotated to one of the six cortical layers according to their correlation coefficient with the six cortical layers in the human brain. Using spatialLIBD (version 1.10.1) ^48^, the correlation between the t-statistics from gene enrichment analysis of histological layers in the reference dataset and the t-statistics from gene enrichment in our query dataset was calculated. The human DLPFC 10x Genomics Visium dataset from Maynard, et al. ^49^ was used for the analysis. The clusters of our query dataset were annotated corresponding to the neuronal layers with the highest correlation coefficients. Layer 1 was annotated based on *RELN* expression and the remaining clusters were assigned to the layers with the highest correlation coefficient scores. Clusters that did not show a positive correlation with any of the six layers in the human brain are designated as ‘undefined’.

#### Developmental trajectory analysis

Partition-based graphical abstraction (PAGA) function, available in the Scanpy library (v1.9.5), within the Python environment (v3.8.18), were employed to create a simplified graph representation of partitions. Utilizing PCA cell embeddings, a neighborhood graph was computed with parameters set to n_pcs = 50 and n_neighbors = 20. Subsequently, the connectivity patterns among partitions was visualized, with edge weights indicating the strength of connections.

#### Transition mapping (TMAP) analysis

TMAP analysis was performed following the methodology as previously described ^50,51^. BrainSpan RNA-seq data ^52^ was used as the primary human brain reference to compare differential gene expression patterns between the human brain during developmental stages and forebrain assembloids. The gene expression patterns of the human brain were derived from ‘brainSpan_pariedVoom_results.rdata’, as utilized in previous studies ^50^. In brief, gene expression levels from RNA-seq data of the human brain tissues were normalized and grouped into 13 stages, ranging from 8 PCW to 40 years. Fold change for each developmental stage was calculated by comparing it to the baseline values of the earliest stage *in vivo* (stage 2; 8-10 PCW) using the limma-voom method ^53^.

Gene expression values from scRNA-seq data of 13 forebrain assembloids were aggregated to generate pseudobulk expression data for each sample. Gene expression values from pseudobulk expression data of 13 forebrain assembloids and bulk RNA-seq data of 3 early brain organoids (d32) were normalized, and fold change was calculated for each of the 13 forebrain assembloids by comparing them to the 3 early brain organoids using DESeq2 (v1.40.2) ^54^. Genes were then ranked by log fold change (logFC), and the rank-rank hypergeometric test^55^ was employed to calculate the significance of the overlap of the genes, using a step size of 200 genes ^51^.

#### Calculation of MI score

Mutual Information (MI) score was calculated based on the previous study ^38^. MI score was calculated between clusters and individual organoids/assembloids with mpmi (v0.43.2.1). The statistical significance of the observed MI scores was calculated by generating background distributions for each dataset.

### Data analysis

Statistical analyses were performed using GraphPad Prism ver. 10. All data were presented as mean +/− SEM. Comparisons between groups were performed using an unpaired *t*-test or a nested *t*-test as described in figure legends. Values of *p* < 0.05 were considered statistically significant.

## Data availability

All data either generated or analyzed during this study are included in this article (and its supplementary information files).

## Code availability

The codes used for data analysis are available upon request.

## Acknowledgements

We thank Mi-Ryoung Song at GIST for their generous provision of the RELN-expressing construct. This research was supported by grants from the National Research Foundation of Korea (NRF-2022R1A2C3002702, RS-2023-00223277), Samsung Science and Technology Foundation (SSTF-BA2101-12), New Faculty Startup Fund from Seoul National University, and the BK21FOUR Research Fellowship.

## Author contributions

E.K, Y.K and K.S conceived the ideas and experimental design. E.K, Y.K, S.H, I.K, and J.L performed the overall experiments. Y.K, J.K, K.Y, and J.C analyzed scRNA-seq data. J.Y performed whole-cell patch-clamp experiments and J.K helped analyze data from electrophysiological experiments. E.K, Y.K and K.S wrote the manuscript.

## Supplementary Figure legends

**Supplementary Fig. 1.**
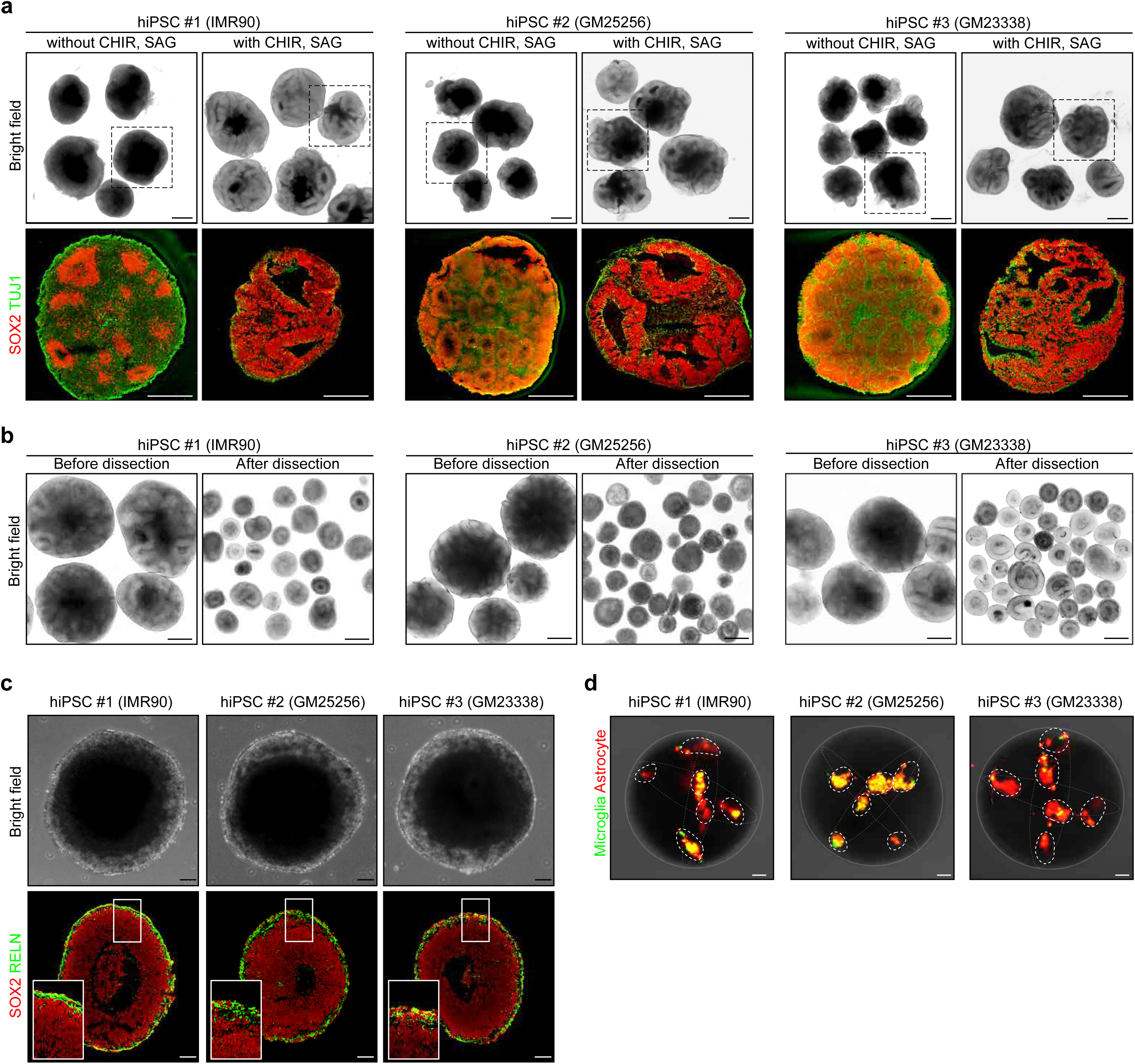
Creation of forebrain assembloids by a module-based, cellular reconstitution of multiple cell types in human brains. (a) (top) Representative bright field images of early forebrain organoids (day 32), derived from hiPSCs, treated with CHIR99021 and SAG. (bottom) Magnified images of forebrain organoids, demarcated by dotted boxes on top panels, immunostained for NPCs (SOX2, red) and neurons (TUJ1, green). Scale bars, 1 mm. (b) Representative bright field images of manually-dissected, single rosettes (left; before dissection, right; after dissection). Scale bars, 1 mm. (c) Representative images of single rosettes encapsulated with the RELN+ layer (day 35) immunostained for SOX2 and RELN. Scale bars, 100 μm. (d) Representative images of intermediate assembloids (day 50) immediately after being microinjected with glial cells. Astrocytes and microglia were labeled with RFP and GFP, respectively. Scale bars, 100 μm.

**Supplementary Fig. 2.**
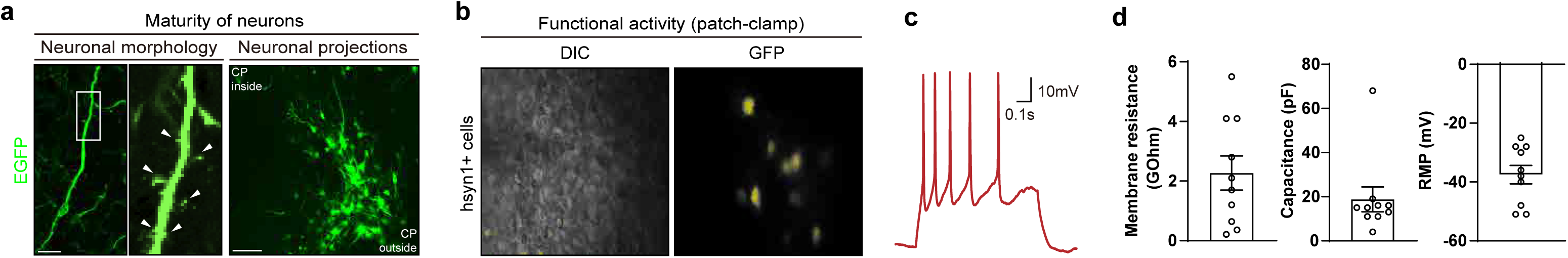
Forebrain assembloids represent neuronal maturity at the single-cell level. (a) (left) Representative images of neurons in forebrain assembloids (d80), showing neuronal morphology. The white arrowheads indicate dendritic spines. Scale bar, 10 μm. (right) Merged images of a series of z sections of forebrain assembloids (d80) with sparsely labeled neurons (EGFP), showing neuronal projections (The sequential scanning video of individual z sections are presented in Supplementary Video 1). CP; cortical plate. Scale bar, 100 μm. (b) Representative images of neurons infected with the AAV-syn1-GFP virus for patch-clamp recording. (c) Representative traces of spontaneous action potential firing in forebrain assembloids. (d) Membrane resistance (left), capacitance (middle), and resting membrane potential (right) of a cell from forebrain assembloids measured using whole-cell patch-clamp recording (Ten cells were recorded; n = 10).

**Supplementary Fig. 3.**
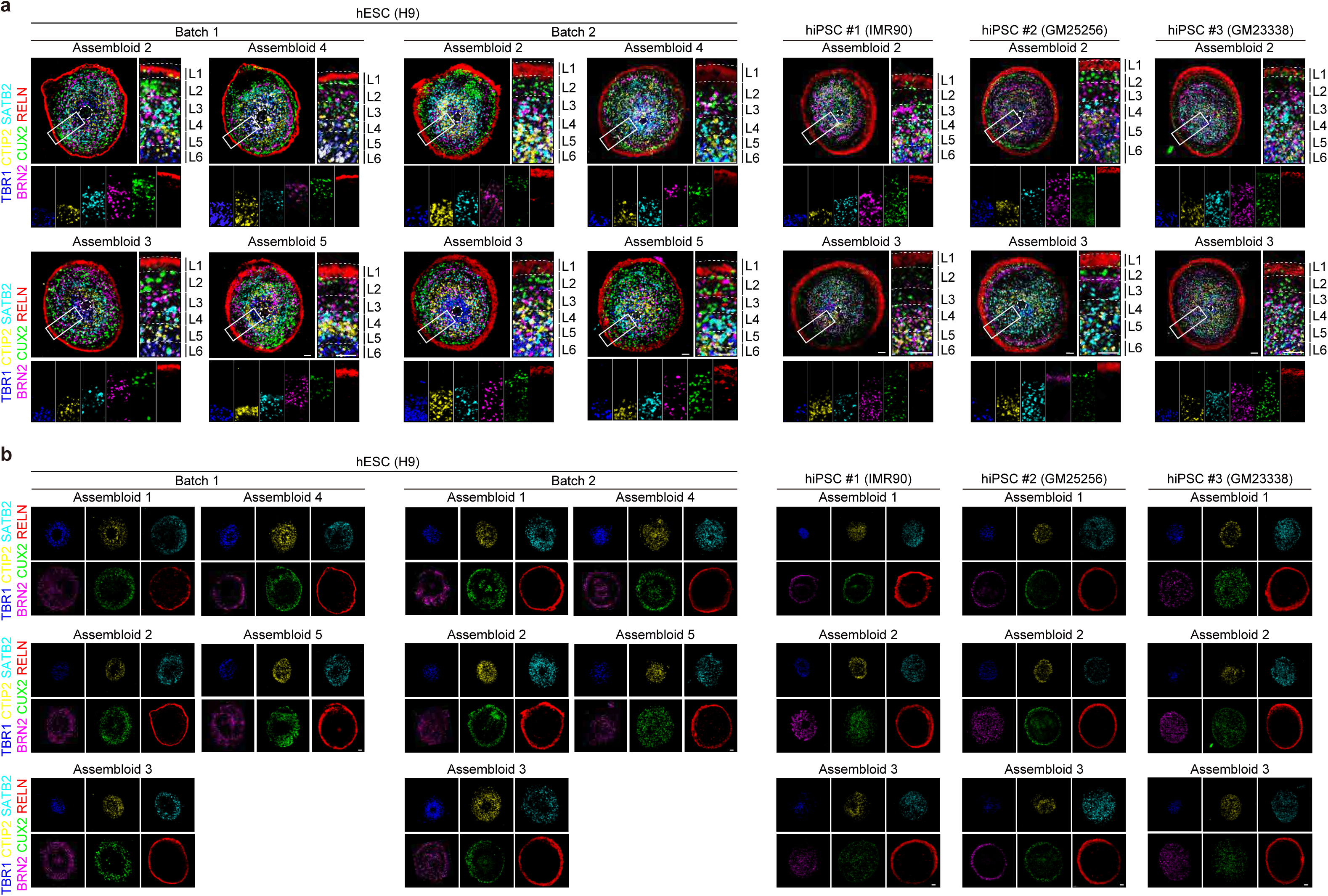
Forebrain assembloids represent the six-layered cortical structure and contain glial cells. (e) Representative images of 6-layered cortical structures of forebrain assembloids (d80). Top left panels show merged images of three serial sections at 8-μm intervals in which each section was immunostained for TBR1/CTIP2, SATB2/RELN, and BRN2/CUX2, respectively. Magnified images (insets in the upper left panels) are shown on the right and bottom panels. Dotted lines demarcate the border of each 6 layer. L1; layer 1, L2; layer 2, L3; layer 3, L4; layer 4, L5; layer 5, L6; layer 6. Scale bars, 100 μm. (b) Images of the TBR1-, CTIP2-, SATB2-, BRN2-, CUX2-, and RELN-positive layers of forebrain assembloids (d80) in Fig. 2c and Supplementary Fig. 3a. Scale bars, 100 μm.

**Supplementary Fig. 4.**
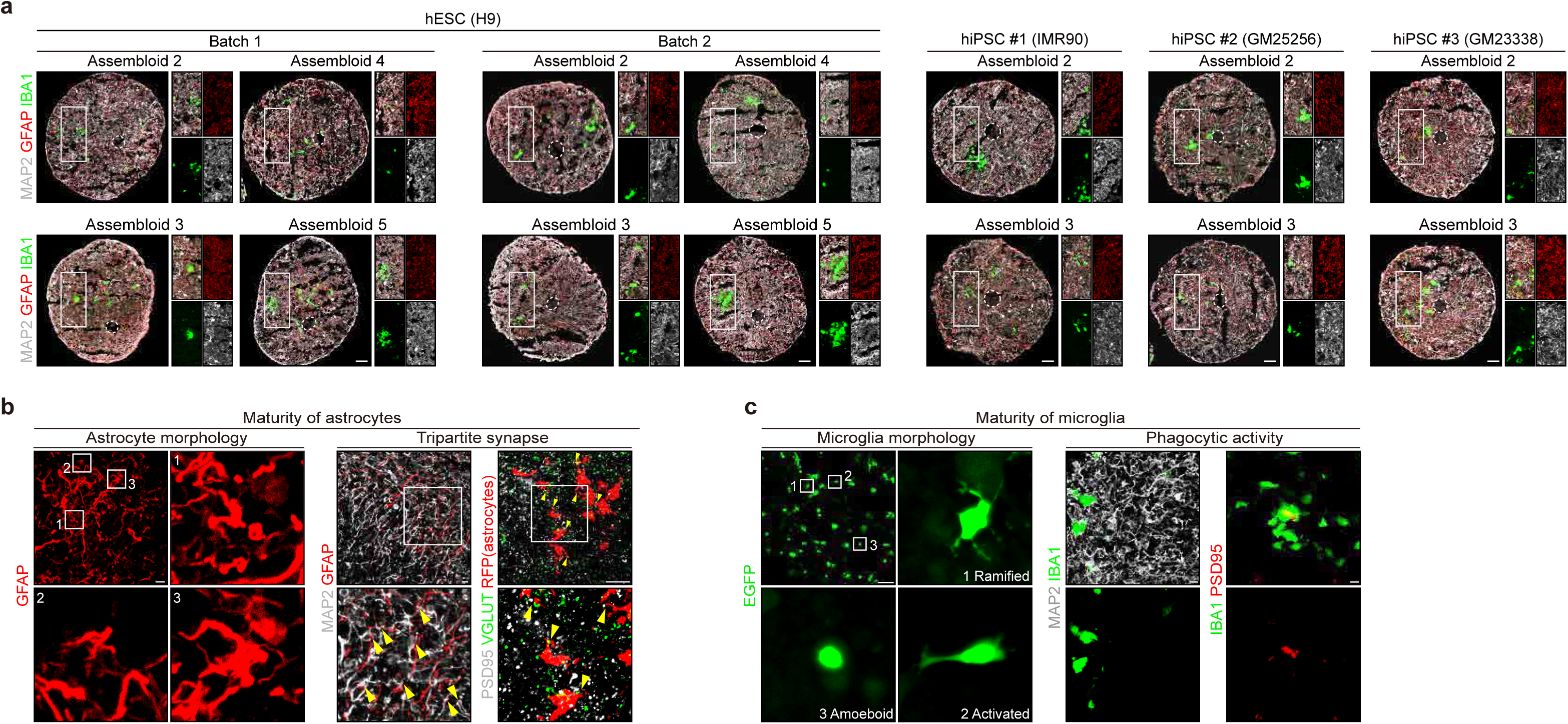
Forebrain assembloids contain GFAP-positive astrocytes and IBA1-positive microglia. (a) Representative images of forebrain assembloids (d80) for neurons (MAP2), astrocytes (GFAP), and microglia (IBA1). Magnified images are shown in the panel on the right. Scale bars, 100 μm. (b) Immunostaining analysis of the morphology and function of astrocytes in forebrain assembloids (d80). Magnified images (insets in the upper panels) are shown below. The yellow arrowheads indicate tripartite synapses. Scale bars, 10 μm. (c) Immunostaining analysis of the morphology and function of microglia in forebrain assembloids (d80). Magnified images for left panels (insets in the upper panel) are shown below. Scale bars, 10 μm.

**Supplementary Fig. 5.**
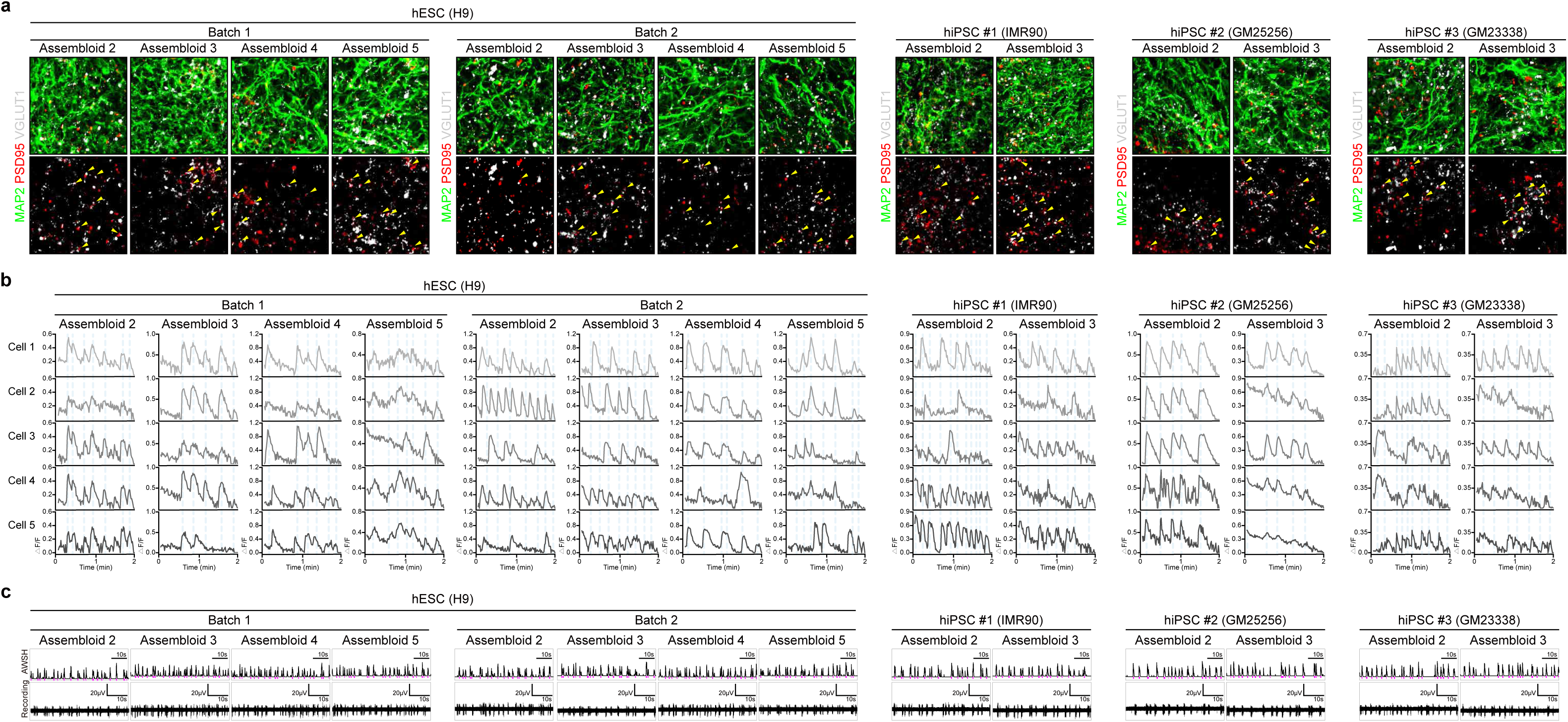
Forebrain assembloids display neuronal activity and functional connectivity. (a) Representative images of forebrain assembloids (d80) immunostained for neurons (MAP2) and synapses (PSD95 and VGLUT1). The yellow arrowheads indicate synapses co-localized with PSD95 and VGLUT1. Scale bars, 50 μm. (b) Representative images of calcium imaging analyses of selected cells in forebrain assembloids (d80). (c) Representative images of the array-wide spike histogram (AWSH) and recording plot analyzed by MEA in forebrain assembloids (d80).

**Supplementary Fig. 6.**
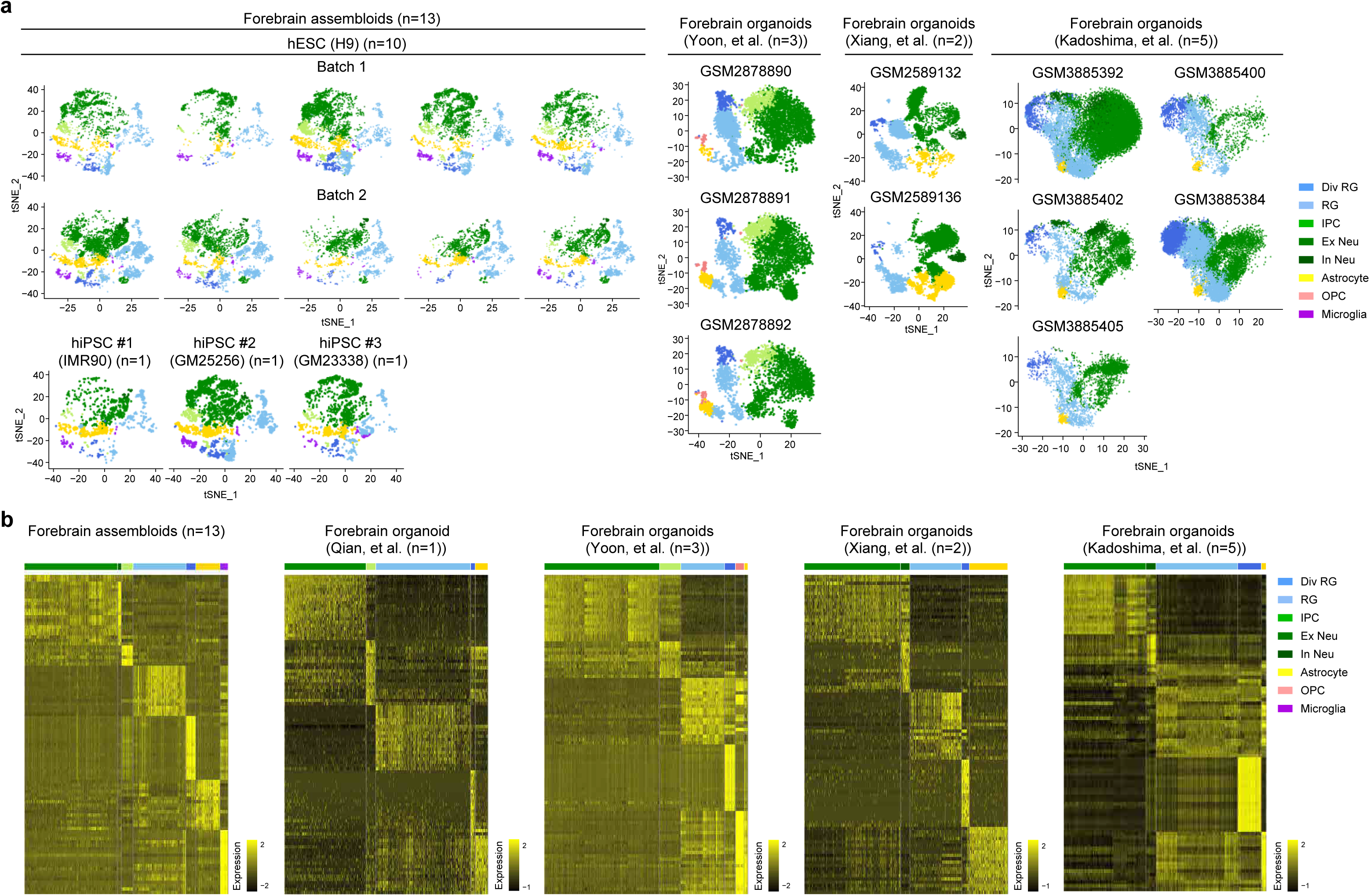
scRNA sequencing analysis for cellular compositions of forebrain assembloids in comparison with current forebrain organoids. (a) tSNE plots of scRNA-seq data from forebrain assembloids (n=13; hESC (H9): five assembloids per batch, two batches analyzed; hiPSC #1 (IMR90), #2 (GM25256), and #3 (GM23338): one assembloid from one batch analyzed) and ten independent, widely-utilized forebrain organoids, developed using methods by Yoon, et al.^33^ (n=3), Xiang, et al.^8^ (n=2), and Kadoshima, et al.^34^ (n=5). Cells are colored by cell type and labeled with cell type annotations. RG, radial glia; Div RG, dividing RG; IPC, intermediate progenitor cell; Ex Neu, excitatory neuron; In Neu, inhibitory neuron; OPC, oligodendrocyte progenitor cell. (b) Heatmap representation of the average gene expression of top 20 marker genes expressed in each cell type in current forebrain organoids and forebrain assembloids.

**Supplementary Fig. 7.**
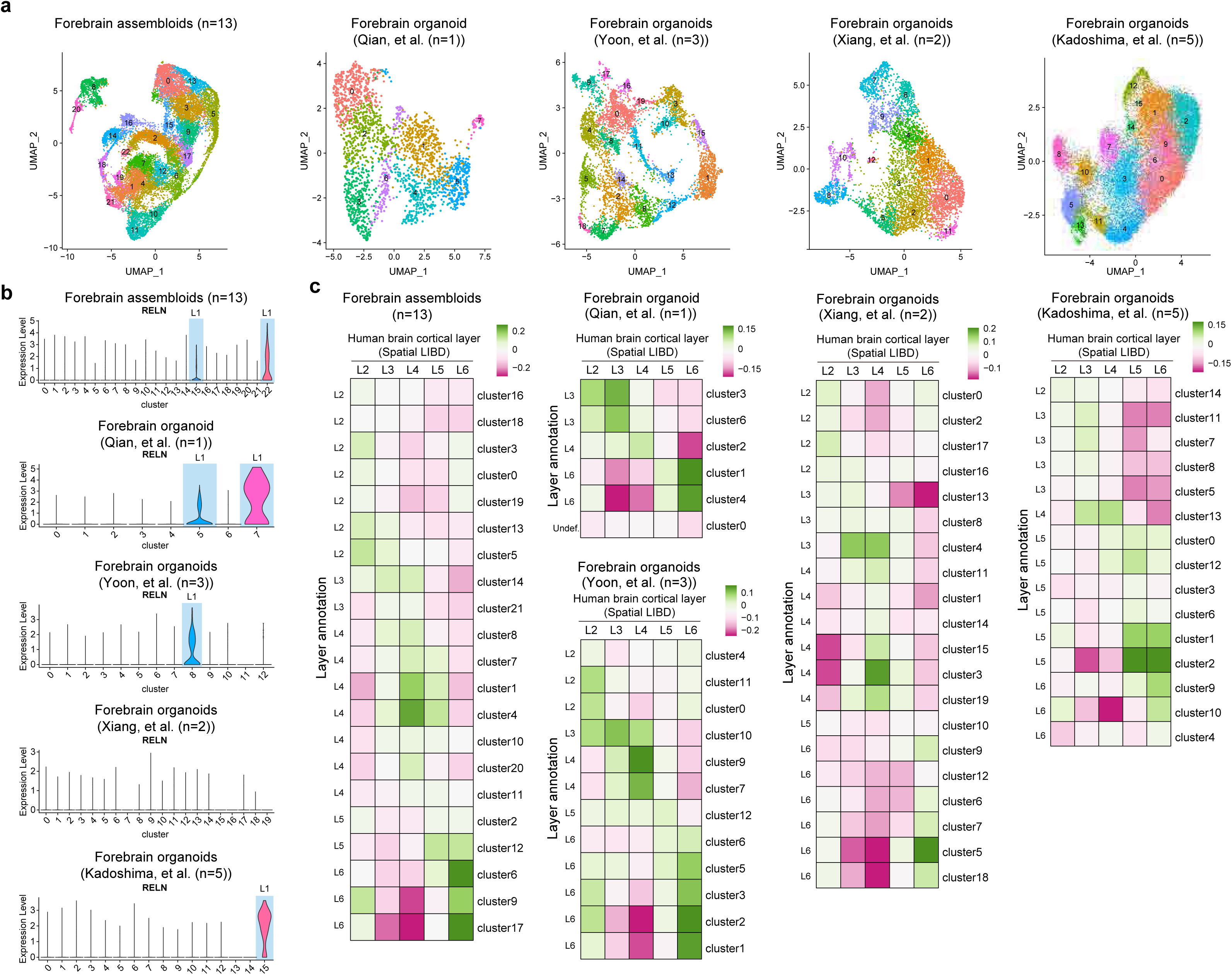
scRNA-seq analysis for cortical layer specification of forebrain assembloids in comparison with current forebrain organoids. (a) UMAP plots of scRNA-seq data of excitatory neurons from forebrain assembloids (n=13; hESC (H9): five assembloids per batch, two batches analyzed; hiPSC #1 (IMR90), #2 (GM25256), and #3 (GM23338): one assembloid from one batch analyzed), and eleven independent, widely-utilized forebrain organoids, developed using methods by Qian, et al.^20^ (n=1), Yoon, et al.^33^ (n=3), Xiang, et al.^8^ (n=2), and Kadoshima, et al.^34^ (n=5). Cells are colored by cell type and labeled with cell type annotations. (b) Violin plots showing the expression of layer 1 marker gene (RELN) in each of excitatory neuron cluster shown in panel ‘a’. Clusters highlighted with light blue color were annotated as layer 1. (c) Heatmaps displaying correlation coefficients between excitatory neuron clusters in forebrain assembloids/organoids, excluding the clusters annotated as layer 1 in (b), and cortical layers (layer 2-6) in the human brain. Excitatory neuron clusters in assembloids/organoids are re-clustered and further annotated to one of the cortical layers according to their correlation coefficient with the cortical layers in the human brain. Clusters that did not show a positive correlation with any of the six layers in the human brain are designated as ‘undefined’. Cells are colored by six cortical layers and labeled with layer annotations.

## Supplementary Video

**Supplementary Video 1. Sequential scanning video of z section images of forebrain assembloids to visualize neuronal projections**

Sparsely labeled neurons with EGFP were visualized in a sequential scanning video of z section images of forebrain assembloids, acquired from confocal microscopy.

**Supplementary Video 2. Live cell imaging analysis for microglia movement in forebrain assembloids**

Movement of EGFP+ microglia in forebrain assembloids was visualized using live cell confocal microscopy.

**Supplementary Video 3. Calcium imaging analysis of forebrain assembloids at bulk area level**

Spontaneous calcium surges in neurons throughout the entire area of forebrain assembloids, represented by Fluo-4-AM fluorescence, were visualized using live cell confocal microscopy.

**Supplementary Video 4. Calcium imaging analysis of forebrain assembloids at single-cell level**

Spontaneous calcium surges in neurons between connected cortical layers within forebrain assembloids, represented by Fluo-4-AM fluorescence, were visualized using live cell confocal microscopy.

## Supplementary Data

**Supplementary Data 1. Summary of scRNA-seq analyses of identified cell clusters in scRNA-seq data of current forebrain organoids and forebrain assembloids**

Per cell type, the overall gene expression, including known marker genes and cell number, analyzed from scRNA-seq data of current forebrain organoids developed by four protocols and forebrain assembloids are listed. Cell types were designated based on the marker genes of each cluster.

## References

1 Lancaster, M. A. et al. Cerebral organoids model human brain development and microcephaly. Nature 501, 373–379 (2013). 10.1038/nature12517

2 Quadrato, G. et al. Cell diversity and network dynamics in photosensitive human brain organoids. Nature 545, 48–53 (2017). 10.1038/nature22047

3 Qian, X. et al. Brain-Region-Specific Organoids Using Mini-bioreactors for Modeling ZIKV Exposure. Cell 165, 1238–1254 (2016). 10.1016/j.cell.2016.04.032

4 Birey, F. et al. Assembly of functionally integrated human forebrain spheroids. Nature 545, 54–59 (2017). 10.1038/nature22330

5 Kadoshima, T. et al. Self-organization of axial polarity, inside-out layer pattern, and species-specific progenitor dynamics in human ES cell&#x2013;derived neocortex. Proceedings of the National Academy of Sciences 110, 20284–20289 (2013). doi:10.1073/pnas.1315710110

6 Jo, J. et al. Midbrain-like Organoids from Human Pluripotent Stem Cells Contain Functional Dopaminergic and Neuromelanin-Producing Neurons. Cell Stem Cell 19, 248–257 (2016). 10.1016/j.stem.2016.07.005

7 Monzel, A. S. et al. Derivation of Human Midbrain-Specific Organoids from Neuroepithelial Stem Cells. Stem Cell Reports 8, 1144–1154 (2017). 10.1016/j.stemcr.2017.03.010

8 Xiang, Y. et al. Fusion of Regionally Specified hPSC-Derived Organoids Models Human Brain Development and Interneuron Migration. Cell Stem Cell 21, 383–398.e387 (2017). 10.1016/j.stem.2017.07.007

9 Bagley, J. A., Reumann, D., Bian, S., Lévi-Strauss, J. & Knoblich, J. A. Fused cerebral organoids model interactions between brain regions. Nature Methods 14, 743–751 (2017). 10.1038/nmeth.4304

10 Andersen, J. et al. Generation of Functional Human 3D Cortico-Motor Assembloids. Cell 183, 1913–1929.e1926 (2020). 10.1016/j.cell.2020.11.017

11 Miura, Y. et al. Generation of human striatal organoids and cortico-striatal assembloids from human pluripotent stem cells. Nature Biotechnology 38, 1421–1430 (2020). 10.1038/s41587-020-00763-w

12 Kelava, I. & Lancaster, Madeline A. Stem Cell Models of Human Brain Development. Cell Stem Cell 18, 736–748 (2016). 10.1016/j.stem.2016.05.022

13 Lancaster, M. A. & Knoblich, J. A. Organogenesis in a dish: Modeling development and disease using organoid technologies. Science 345, 1247125 (2014). doi:10.1126/science.1247125

14 Quadrato, G., Brown, J. & Arlotta, P. The promises and challenges of human brain organoids as models of neuropsychiatric disease. Nature Medicine 22, 1220–1228 (2016). 10.1038/nm.4214

15 Stiles, J. & Jernigan, T. L. The Basics of Brain Development. Neuropsychology Review 20, 327–348 (2010). 10.1007/s11065-010-9148-4

16 Quadrato, G. & Arlotta, P. Present and future of modeling human brain development in 3D organoids. Current Opinion in Cell Biology 49, 47–52 (2017). 10.1016/j.ceb.2017.11.010

17 Causeret, F., Moreau, M. X., Pierani, A. & Blanquie, O. The multiple facets of Cajal-Retzius neurons. Development 148 (2021). 10.1242/dev.199409

18 Bystron, I., Blakemore, C. & Rakic, P. Development of the human cerebral cortex: Boulder Committee revisited. Nature Reviews Neuroscience 9, 110–122 (2008). 10.1038/nrn2252

19 Giandomenico, S. L. et al. Cerebral organoids at the air–liquid interface generate diverse nerve tracts with functional output. Nature Neuroscience 22, 669–679 (2019). 10.1038/s41593-019-0350-2

20 Qian, X. et al. Sliced Human Cortical Organoids for Modeling Distinct Cortical Layer Formation. Cell Stem Cell 26, 766–781.e769 (2020). 10.1016/j.stem.2020.02.002

21 Xu, R. et al. Developing human pluripotent stem cell-based cerebral organoids with a controllable microglia ratio for modeling brain development and pathology. Stem Cell Reports 16, 1923–1937 (2021). 10.1016/j.stemcr.2021.06.011

22 Sloan, S. A. et al. Human Astrocyte Maturation Captured in 3D Cerebral Cortical Spheroids Derived from Pluripotent Stem Cells. Neuron 95, 779–790.e776 (2017). 10.1016/j.neuron.2017.07.035

23 Kim, E. et al. Creation of bladder assembloids mimicking tissue regeneration and cancer. Nature 588, 664–669 (2020). 10.1038/s41586-020-3034-x

24 Neumann, J. E. et al. A mouse model for embryonal tumors with multilayered rosettes uncovers the therapeutic potential of Sonic-hedgehog inhibitors. Nat Med 23, 1191–1202 (2017). 10.1038/nm.4402

25 Wang, L., Hou, S. & Han, Y.-G. Hedgehog signaling promotes basal progenitor expansion and the growth and folding of the neocortex. Nature Neuroscience 19, 888–896 (2016). 10.1038/nn.4307

26 Muffat, J. et al. Efficient derivation of microglia-like cells from human pluripotent stem cells. Nature Medicine 22, 1358–1367 (2016). 10.1038/nm.4189

27 Tcw, J. et al. An Efficient Platform for Astrocyte Differentiation from Human Induced Pluripotent Stem Cells. Stem Cell Reports 9, 600–614 (2017). 10.1016/j.stemcr.2017.06.018

28 Saito, T. et al. Neocortical Layer Formation of Human Developing Brains and Lissencephalies: Consideration of Layer-Specific Marker Expression. Cerebral Cortex 21, 588–596 (2010). 10.1093/cercor/bhq125

29 Hevner, R. F. Layer-Specific Markers as Probes for Neuron Type Identity in Human Neocortex and Malformations of Cortical Development. Journal of Neuropathology & Experimental Neurology 66, 101–109 (2007). 10.1097/nen.0b013e3180301c06

30 Buzsáki, G. Large-scale recording of neuronal ensembles. Nature Neuroscience 7, 446–451 (2004). 10.1038/nn1233

31 Tolonen, M., Palva, J. M., Andersson, S. & Vanhatalo, S. Development of the spontaneous activity transients and ongoing cortical activity in human preterm babies. Neuroscience 145, 997–1006 (2007). 10.1016/j.neuroscience.2006.12.070

32 Fries, P. A mechanism for cognitive dynamics: neuronal communication through neuronal coherence. Trends in Cognitive Sciences 9, 474–480 (2005). 10.1016/j.tics.2005.08.011

33 Yoon, S.-J. et al. Reliability of human cortical organoid generation. Nature Methods 16, 75–78 (2019). 10.1038/s41592-018-0255-0

34 Kadoshima, T. et al. Self-organization of axial polarity, inside-out layer pattern, and species-specific progenitor dynamics in human ES cell-derived neocortex. Proc Natl Acad Sci U S A 110, 20284–20289 (2013). 10.1073/pnas.1315710110

35 Lake, B. B. et al. Integrative single-cell analysis of transcriptional and epigenetic states in the human adult brain. Nature Biotechnology 36, 70–80 (2018). 10.1038/nbt.4038

36 Lake, B. B. et al. Neuronal subtypes and diversity revealed by single-nucleus RNA sequencing of the human brain. Science 352, 1586–1590 (2016). doi:10.1126/science.aaf1204

37 Cadwell, C. R., Bhaduri, A., Mostajo-Radji, M. A., Keefe, M. G. & Nowakowski, T. J. Development and Arealization of the Cerebral Cortex. Neuron 103, 980–1004 (2019). 10.1016/j.neuron.2019.07.009

38 Velasco, S. et al. Individual brain organoids reproducibly form cell diversity of the human cerebral cortex. Nature 570, 523–527 (2019). 10.1038/s41586-019-1289-x

39 Dulabon, L. et al. Reelin Binds &#x3b1;3&#x3b2;1 Integrin and Inhibits Neuronal Migration. Neuron 27, 33–44 (2000). 10.1016/S0896-6273(00)00007-6

40 Tidball, A. M. et al. Deriving early single-rosette brain organoids from human pluripotent stem cells. Stem Cell Reports 18, 2498–2514 (2023). 10.1016/j.stemcr.2023.10.020

41 Wang, Y. et al. Modeling human telencephalic development and autism-associated SHANK3 deficiency using organoids generated from single neural rosettes. Nature Communications 13, 5688 (2022). 10.1038/s41467-022-33364-z

42 Cao, S. Y. et al. Enhanced derivation of human pluripotent stem cell-derived cortical glutamatergic neurons by a small molecule. Sci Rep 7, 3282 (2017). 10.1038/s41598-017-03519-w

43 Zheng, G. X. Y. et al. Massively parallel digital transcriptional profiling of single cells. Nature Communications 8, 14049 (2017). 10.1038/ncomms14049

44 Hao, Y. et al. Integrated analysis of multimodal single-cell data. Cell 184, 3573–3587.e3529 (2021). 10.1016/j.cell.2021.04.048

45 Germain, P. L., Lun, A., Garcia Meixide, C., Macnair, W. & Robinson, M. D. Doublet identification in single-cell sequencing data using scDblFinder. F1000Res 10, 979 (2021). 10.12688/f1000research.73600.2

46 Haghverdi, L., Lun, A. T. L., Morgan, M. D. & Marioni, J. C. Batch effects in single-cell RNA-sequencing data are corrected by matching mutual nearest neighbors. Nat Biotechnol 36, 421–427 (2018). 10.1038/nbt.4091

47 Bhaduri, A. et al. Cell stress in cortical organoids impairs molecular subtype specification. Nature 578, 142–148 (2020). 10.1038/s41586-020-1962-0

48 Pardo, B. et al. spatialLIBD: an R/Bioconductor package to visualize spatially-resolved transcriptomics data. BMC Genomics 23, 434 (2022). 10.1186/s12864-022-08601-w

49 Maynard, K. R. et al. Transcriptome-scale spatial gene expression in the human dorsolateral prefrontal cortex. Nature Neuroscience 24, 425–436 (2021). 10.1038/s41593-020-00787-0

50 Gordon, A. et al. Long-term maturation of human cortical organoids matches key early postnatal transitions. Nature Neuroscience 24, 331–342 (2021). 10.1038/s41593-021-00802-y

51 Stein, Jason L. et al. A Quantitative Framework to Evaluate Modeling of Cortical Development by Neural Stem Cells. Neuron 83, 69–86 (2014). 10.1016/j.neuron.2014.05.035

52 Li, M. et al. Integrative functional genomic analysis of human brain development and neuropsychiatric risks. Science 362, eaat7615 (2018). doi:10.1126/science.aat7615

53 Ritchie, M. E. et al. limma powers differential expression analyses for RNA-sequencing and microarray studies. Nucleic Acids Research 43, e47–e47 (2015). 10.1093/nar/gkv007

54 Love, M. I., Huber, W. & Anders, S. Moderated estimation of fold change and dispersion for RNA-seq data with DESeq2. Genome Biol 15, 550 (2014). 10.1186/s13059-014-0550-8

55 Plaisier, S. B., Taschereau, R., Wong, J. A. & Graeber, T. G. Rank–rank hypergeometric overlap: identification of statistically significant overlap between gene-expression signatures. Nucleic Acids Research 38, e169–e169 (2010). 10.1093/nar/gkq636

